# ELEN – Predicting Loop Quality in Protein Structure Models

**DOI:** 10.1101/2025.08.07.668890

**Authors:** Florian Wieser, Sigrid Kaltenbrunner, Andreas Zechner, Martin Zach, Thomas Pock, Gustav Oberdorfer

## Abstract

Typically, sequences designed *de novo* are assessed *in silico* using deep learning-based protein structure prediction methods prior to wetlab testing. While these deep learning (DL) models excel at predicting well-ordered regions, accurate prediction of loop regions, which often are flexible and crucial for protein function, remains a significant challenge. To address this, we introduce the Equivariant Loop Evaluation Network (ELEN), a local model quality assessment (MQA) method that is tailored towards evaluating the accuracy of protein loops at the per-residue level. ELEN jointly predicts three quality metrics, local Distance Difference Test (lDDT), Contact Area Difference Score (CAD-score), and Root Mean Squared Deviation (RMSD), by comparing predicted to experimental reference structures. Learning these metrics simultaneously enables ELEN to capture complementary structural insights, providing a richer assessment of model accuracy. The network operates at all-atom resolution and employs 3D equivariant group convolutions to learn the local geometric environment of each atom. By incorporating sequence embeddings from large language models (LLMs), such as SaProt, we enhance the sequence and evolutionary awareness of the model. Furthermore, by informing ELEN with per-residue physicochemical features, the model achieves competitive accuracy relative to state-of-the-art MQA methods on the Continuous Automated Model EvaluatiOn (CAMEO) benchmark. Although ELEN was primarily developed for assessing loop quality, its architecture also demonstrates strong potential for general MQA tasks. We used ELEN to perform detailed analysis, including identification of flexible or disordered regions and assessment of structural effects from single-residue mutations on three sets of redesigned enzymes. We show that for all sets ELEN successfully identifies poor design positions and thus serves as a powerful tool for advancing both the study and modeling of loops in protein structures.

## 2. Introduction

Proteins are central to virtually all cellular processes, yet experimental structure determination for every novel or designed protein remains prohibitively time-consuming, expensive, or may even fail entirely. As a result, computational structure prediction has become a foundation of modern protein science and engineering. Driven largely by advances in DL, recent prediction methods have approached the resolution of experimental techniques and are expected to improve further in the coming years [1]. Current prominent examples include AlphaFold2 (AF2)[2], AlphaFold3 (AF3) [3], RosettaFold2 [4], and the Evolutionary Scale Model family (ESM-2 [5] and ESM-3 [6]. Despite these advances, predicted protein models still contain errors that can limit their utility for answering biologically relevant questions. This is especially true for proteins that lack critical information, such as homologous structures as templates or sequences for residue-residue coevolution data in their multiple sequence alignments (MSAs) [7]. A persistent challenge is the accurate modeling of loop regions: prediction accuracy decreases with increasing loop length, the frequency of loop residues tends to be underpredicted [8] and many loops are inherently disordered.

Although the internal confidence metric of AF2, predicted lDDT (plDDT) is a valuable quality indicator, orthogonal MQA methods remain important for identifying local errors (local MQA) and selecting optimal models (global MQA) [9] [10] [11]. MQA approaches are typically categorized as single-model or multi-model methods. The former evaluate each structure independently and are computationally efficient, while the latter leverage ensembles but are more resource-intensive [12]. Notably, single-model methods have recently surpassed multi-model methods in accuracy, as demonstrated in the 14th Critical Assessment of protein Structure Prediction (CASP14) [11]. Whereas early MQA relied on knowledge-based energy functions, modern methods increasingly use machine learning (ML) and DL techniques, leveraging diverse architectures and features. Early approaches such as the ProQ and ModFOLD families relied on traditional ML models that combined multiple sequence- and structure-derived features [10] [11] [12] [13] [14] [15] [16]. Both families have since evolved to incorporate DL architectures and consensus-based strategies, which has led to significant improvements in accuracy. The ModFOLD family currently represents the state-of-the-art among MQA methods, consistently ranking at the top in independent blind evaluations by integrating outputs from several methods using neural networks [11] [16].

A significant advance in the field has been the introduction of 3D-equivariant neural network architectures, such as EnQA [7] and EDN [17]. Based on the principles of Geometric Deep Learning (GDL), these models efficiently incorporate and process symmetry information [40]. Equivariance ensures that transformations such as rotations and translations applied to the input result in predictable transformations of the output, while invariance is a special case where the output remains unchanged regardless of input transformation [18]. For MQA, invariance at the output (model-level invariance) is desirable to ensure that quality assessments are independent of a model’s orientation or position, while equivariant layers enable the network to accurately detect and analyze structural motifs regardless of their orientation [19] [17]. Motivated by the simplicity and effectiveness of the Equivariant Deep Network (EDN) architecture [17], we extend and modify this framework to specifically assess the quality of loop regions on a per-residue basis. Key modifications and extensions include the focused training specifically on loop pockets extracted from AF2-predicted protein structures, the shifting of the prediction target from global quality to detailed per-residue quality scores and the simultaneous use of multiple complementary per-residue structural quality metrics. In addition, we integrate physicochemical and sequence-based features, which further boosts the ability of ELEN to comprehensively and accurately assess loop quality.

Collectively, these enhancements form the Equivariant Loop Evaluation Network, which, to the best of our knowledge, constitutes the first MQA method that is specifically tailored to assessing the quality of loop residues. ELEN’s per-residue error predictions offer critical insights into local structural confidence that support detailed model analysis and targeted refinement. By aggregating these per-residue scores, ELEN produces a robust global metric for loop quality, and thereby enables the reliable evaluation of the overall structural accuracy and the effective ranking of candidate loop models to identify those that most closely resemble native conformations. Benchmarking against ground truth structural deviations reveals that the performance of ELEN is on par with, or exceeds, that of leading consensus methods (see Results). By leveraging high-level structural representations, including features like hydrogen bond counts and sequence embeddings from LLMs, ELEN accurately identifies structural inaccuracies, loop flexibility, and missing electron densities in crystal structures. Importantly, ELEN’s strong performance is not restricted to loop residues, but extends to all types of secondary structure elements. Its flexible architecture further enables the assessment of a broad range of structural and functional regions within protein models. We show that practical applicability of ELEN by using it to asses the model quality of loops in a set of redesigned loops in a naturally occurring enzyme as well as its ability to identify sub optimal backbone - side chain pairs in *de novo* designed retro-aldolases and to interpret reasons for partial misfolding in a design kemp eliminase.

## 3. Methods

### 3.1 Neural network architecture

Our model builds upon the equivariant convolutional neural network framework EDN introduced by Eismann et al. in [17], which was originally designed for the prediction of a global quality score for entire protein structures using only 3D atomic coordinates and their respective element types. Unlike conventional 3D convolutional neural networks that operate on voxelized grids, EDN employs a point cloud representation where convolutional operations act upon features assigned to individual atoms or residues within local neighborhoods. A key property of this architecture is its rotation equivariance, which ensures that rotations of the input structure result in corresponding rotations of the network’s internal representations. This enables the network to capture molecular symmetries without the need for extensive rotational data augmentation. Rotation equivariance is achieved by defining convolutional filters as truncated series expansions of spherical harmonics with learnable radial functions. These filters detect local structural motifs irrespective of their absolute orientation or spatial position, while preserving relative spatial arrangements and orientations.

We build on this architecture and extend the network’s output from a single global score to detailed per-residue quality predictions. Specifically, ELEN predicts three distinct residue-level metrics: lDDT [20], CAD-score [21], and RMSD [39]. Each metric addresses unique aspects of structural quality, and their combination provides a comprehensive and nuanced assessment. RMSD quantifies the average atomic distances after structural superposition and is widely adopted for its intuitive interpretation. However, it is sensitive to outliers and structure size, and requires structural alignment. In contrast, lDDT and CAD-score are superposition-free, contact-based metrics. The lDDT metric evaluates local atomic distance deviations and serves as the primary accuracy criterion in CASP (EMA category) [1] and CAMEO [23]. The CAD-score measures differences in residue contact surface areas and strongly correlates with physical realism metrics such as MolProbity [22]. While lDDT and CAD-score are highly correlated (Spearman’s ρ = 0.91), we include RMSD because it shows lower correlations with lDDT (ρ = 0.76) and CAD-score (ρ = 0.69).

This allows the network to capture additional, orthogonal aspects of structural quality and improves sensitivity and specificity. Because RMSD is inversely correlated with model quality and operates on a different scale than lDDT and CAD-score—which both naturally lie in the range [0, 1]—RMSD predictions are normalized by the maximum RMSD observed during training and subsequently inverted to ensure they are directionally consistent with the other metrics.

The combined ELEN score of any residue x is computed as:

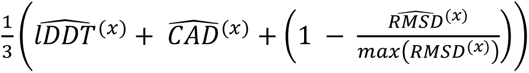

However, each quality metric can also be used individually or combined, providing flexible analysis options.

Additionally, ELEN integrates physicochemical features at the atomic and residue levels. Initially, atomic representations are encoded using one-hot vectors of element types and subsequently enriched by projecting residue-level physicochemical properties (Table 1) onto each respective atom. After pooling atomic representations at the alpha-carbon level, these residue-level physicochemical properties are added again to further refine the learned residue-level representations. Furthermore, the layer sizes of both 3D convolutional and the final dense layers have been increased relative to the original architecture [17], thus enhancing the network’s representational capacity. The architecture is depicted in Figure 1C.

**Figure 1.**
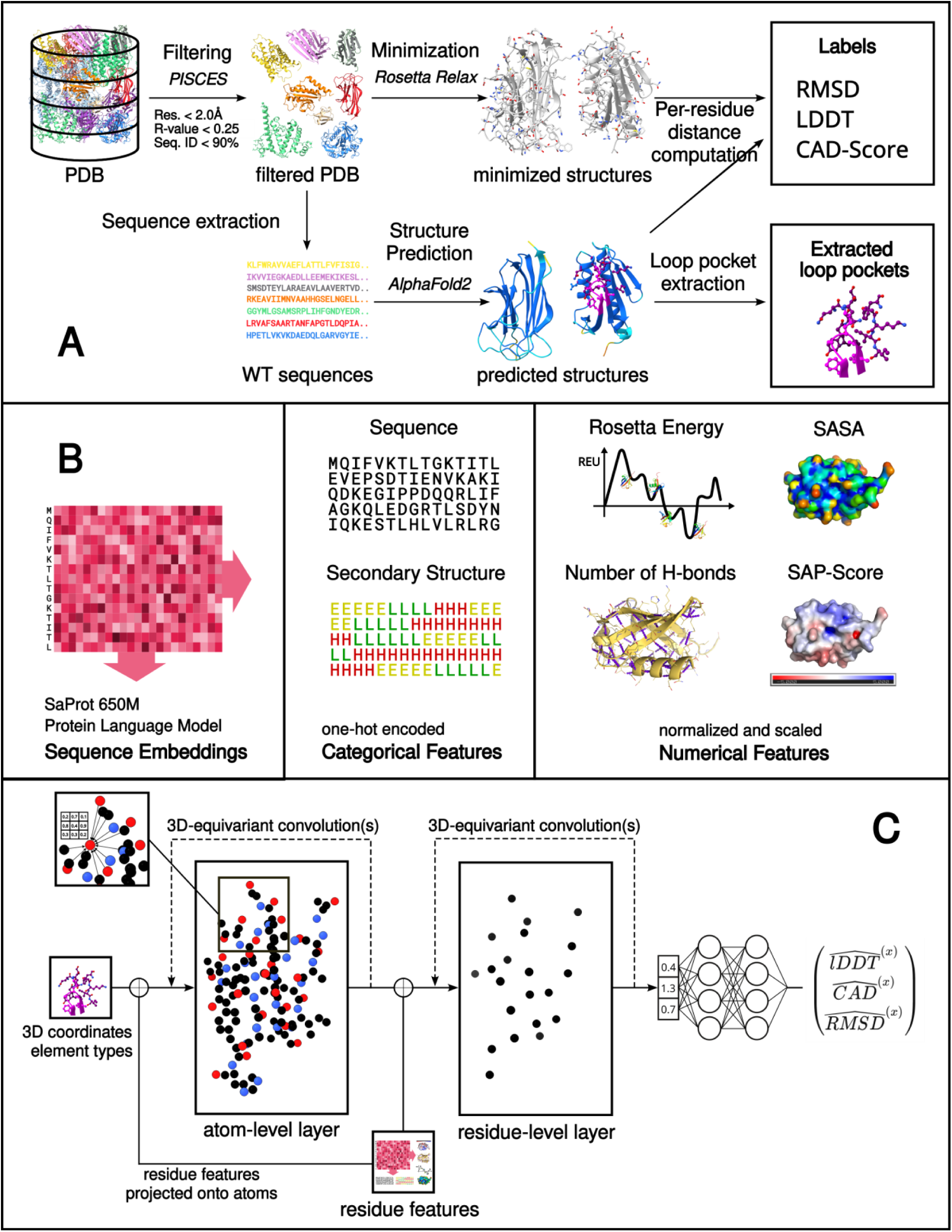
Overview of the ELEN model training pipeline. (A) Data Preparation: Flowchart outlining each step in preparing data for the ELEN network, beginning with protein structure selection, followed by computational preprocessing and coordinate extraction. (B) Feature Generation: Illustration of per-residue feature extraction. Sequence embeddings are obtained from the pretrained SaProt 650M protein language model. Categorical features, including amino acid sequences and predicted secondary structures, are encoded via one-hot encoding. Numerical features, such as Rosetta energy scores, SASA, the number of hydrogen bonds and SAP-score are normalized and scaled prior to their use in training. (C) Network Architecture: Schematic representation of the ELEN architecture. The network initially processes atomic coordinates and their elemental identities through multiple iterations of 3D equivariant convolutions, enabling atoms to capture local geometric environments. Atomic features are subsequently aggregated at the residue-level (C-alpha carbons) and further processed by additional equivariant convolutions. Additionally, initial geometric representations at atomic and residue levels are then integrated with precomputed features from panel B. Finally, aggregated residue-level features are averaged and passed through dense neural network layers to predict residue-specific quality scores (y(l)), which are compared against the ground truth labels during training.

**Table 1.** Overview of input features and their inclusion in ELEN model variants. Summary of all input features provided to the ELEN network, including their type, hierarchical level (atom or residue), calculation source, and encoding format. The table also indicates which features are included in the main ELEN model and in each ablation variant (ELEN-NoSeq, ELEN-GeomOnly).

| Feature Name | Type | Level | Source/Methods | One-hot/Numerical | ELEN | ELEN-NoSeq | ELEN-GeomOnly | Comments |
| --- | --- | --- | --- | --- | --- | --- | --- | --- |
| Atomic coordinates | Structural | Atom | PDB | Numerical | ✓ | ✓ | ✓ | always included |
| Element type | Chemical | Atom | PDB | One-hot | ✓ | ✓ | ✓ | always included |
| Amino acid identity | Sequence | Residue | PDB | One-hot | ✓ | ✓ |  | 20 standard amino acids |
| Secondary Structure | Structural | Residue | DSSP | One-hot | ✓ | ✓ |  | H/E/L 3 state |
| Rosetta energies | Energy | Residue | RosettaScripts | Numerical | ✓ | ✓ |  | Per-residue Rosetta terms |
| Hydrogen bonds | Structural | Residue | RosettaScripts | Numerical | ✓ | ✓ |  | Number of H-bonds |
| SAP score | Structural | Residue | RosettaScripts | Numerical | ✓ | ✓ |  | Spatial aggregation propensity |
| SASA | Structural | Residue | RosettaScripts | Numerical | ✓ | ✓ |  | Solvent-accessible surface area |
| SaProt embedding | Sequence | Residue | SaProt | Numerical | ✓ |  |  | Obtained from the 650M_PDB model |
| Epochs | - | - | - | - | 10 | 32 | 47 | - |

### 3.2 Dataset preparation

#### 3.2.1 Training dataset preparation

To curate a non-redundant training dataset (Figure 1A), we used the Protein Sequence Culling Server (PISCES) [24] to filter a snapshot of the Protein Data Bank (PDB) [25] dated September 2023. We filtered for high structural quality and sequence diversity, by specifically enforcing a maximum sequence identity threshold of 90%, a resolution better than or equal to 2.0 Å and a maximum R-value of 0.25. Subsequently, the resulting 17956 protein crystal structures have been cleaned and minimized using Rosetta FastRelax [41]. The refined structures were then separated into individual protein chains. For each chain, we generated one decoy model using AF2 implemented through LocalColabFold, a local adaptation of the ColabFold pipeline [26]. Unstructured regions were identified based on hydrogen bonding using the Definition Secondary Structure of Protein (DSSP) [27] algorithm. We extracted loops that consist of 2–10 residues that are flanked on both sides by at least 4 residues in secondary structure elements (helix or sheet), as assigned by DSSP. Following the approach in [26], these loops were extracted as ‘loop pockets’, encompassing not only the loop residues themselves but also the surrounding residues, thereby capturing structural context (see Figure 2D for an example). Our loop pockets include all the loop residues themselves, along with the n nearest neighboring residues from the central residue of the loop, extending up to 40 residues in total, to obtain a consistent batch size.

**Figure 2.**
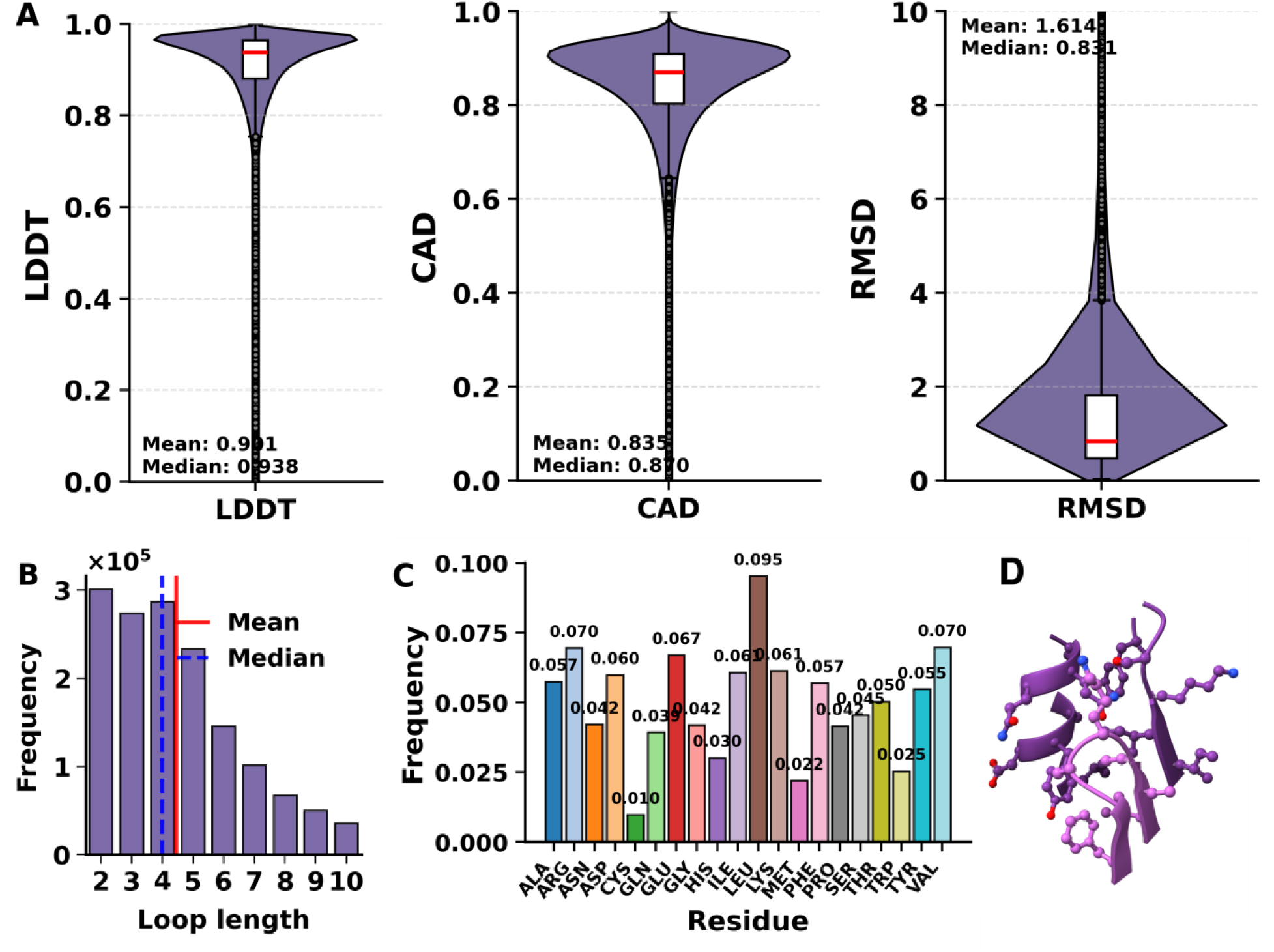
Summary statistics and visualization of the ELEN training dataset. (A) Distributions of per-residue lDDT, CAD-score, and RMSD for all extracted loop pockets, with mean and median values indicated. (B) Histogram of loop lengths for all loops in the dataset (n ≈ 1 million), showing loop length frequency, mean and median. (C) Amino acid composition of extracted loops, normalized by frequency. (D) Example structure of an extracted loop pocket (magenta), illustrating the inclusion of both the loop (light) and its local structural context (dark) used as input for model training.

We computed per-residue distance-based metrics—lDDT, CAD-score, and RMSD—between each AF2 model and its corresponding relaxed crystal structure, ensuring the structures were properly aligned by performing these calculations prior to loop extraction. These metrics serve as the ground-truth labels for model training. Specifically, lDDT [20] was computed using the standalone executable version 1.2. CAD-score [21] was calculated utilizing Voronota software, version 1.27.3834. RMSD was determined using RosettaScripts [28], employing the all-atom variant of RMSD calculation. When splitting the data into training, validation, and testing sets, attention was given to ensure that each PDB identifier (e.g. 1UBQ) appeared in only one set to prevent data leakage. The resulting training, validation and test datasets comprised 938000, 186000 and 18500 loop pockets, respectively.

#### 3.2.2 Benchmark dataset preparation

We test the local and global scoring capabilities of ELEN on a dataset that was obtained from the CAMEO online server [23] in the category of Model Quality Estimation (MQE) in March 2025 and covers a time period of the past three months. Participating methods were ModFOLD7_lDDT [14], ModFOLD8 [11], ModFOLD9 [16], ProQ3 [29], ProQ3D [30], ProQ3D_lDDT [30], QMEANDisCo_3 [31], QMEAN_3 [32], VoroMQA_sw5 and VoroMQA_v2 [33]. Methods that did not provide scores for all models were excluded from the analysis. The final dataset comprises 952 protein models that correspond to 98 protein targets and result in 8681 extracted loops (identified as described in the training dataset preparation) and about 246,000 individual residues. Performance metrics (Pearson and Spearman correlations) were computed with respect to the ground-truth lDDT values provided by CAMEO. Scoring was performed either on loop residues alone or on all residues, allowing both local (loop-focused) and global evaluations.

### 3.3 Feature generation and integration

To provide a richer local representation, ELEN integrates both structural and sequence-derived descriptors at the residue level. These per-residue descriptors are either used directly at the residue level or, when necessary, projected uniformly onto all atoms within the residue to ensure compatibility with atom-level convolution operations. Numerical structural features comprise the number of hydrogen bonds, Rosetta-derived energy terms, Spatial Aggregation Propensity (SAP) scores, and Solvent-Accessible Surface Area (SASA), all precomputed using RosettaScripts [28]. One-hot encoded features comprise the amino acid identity (covering the 20 standard amino acids) and secondary structure types (helix, sheet, or loop) as identified by DSSP. ELEN additionally incorporates sequence-derived embeddings from large protein language models. We selected the SaProt_650M_PDB model [34], which yielded the best accuracy across our benchmarks. Other variants, including embeddings from ESM-2 and smaller SaProt models, were tested but not included in the final model. Before concatenation, features are scaled to ensure numerical stability and effective training. Atom-level features are scaled by a factor of 0.5, and residue-level features are scaled by a factor of 10. We evaluated several normalization strategies, including min-max normalization, mean centralization, and standardization (Z-score), but found that simple scaling without further normalization consistently yielded the best predictive performance. Finally, all features are concatenated either to the initial atomic embedding or, in the case of residue-level features, to the learned residue-level representation.

To evaluate the contribution of different feature types, we developed two ablation variants of the model: ELEN-NoSeq, which excludes sequence embeddings, and ELEN-GeomOnly, which uses only atomic coordinates and element types. These variants help isolate the impact of sequence-derived and higher-level structural features. Notably, we observed that models with richer feature sets required fewer training epochs to reach optimal performance, while more limited models benefited from additional training (see Table 1). A visual overview of the input features and their hierarchical integration into the network architecture is shown in Figure 1B. All feature ablation experiments were performed using a randomly sampled subset of the ELEN loop dataset that contained approximately 100,000 extracted loop pockets, with models trained for a single epoch to efficiently screen feature utility. Predictive performance was quantified using Pearson correlation and mean absolute error (MAE) between model predictions and ground truth labels (lDDT, CAD-score, RMSD).

### 3.4 Implementation details

The final ELEN model comprises approximately 341,000 trainable parameters and was implemented in PyTorch Lightning. Training was performed for ten epochs on an NVIDIA A100 GPU (40 GB RAM) with a batch size of 32 and a learning rate of 1×10⁻⁴, using the Adam optimizer. We used the Huber loss function to balance sensitivity to small errors with robustness to outliers. The total Huber loss was computed across all three per-residue quality metrics (lDDT, CAD-score, and RMSD) and defined as:

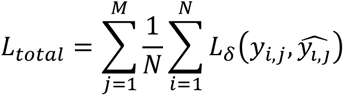

Where N is the number of residues in the batch, M=3 (the number of labels), 𝑦_𝑖,𝑗_ and 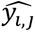 are the true and predicted values for residue i and metric j, and L_total_ is the Huber loss with threshold δ (set to 1.0). All code and training pipelines were developed using Python 3.10, PyTorch 2.x, and associated scientific computing libraries.

### 3.5 Experimental evaluation of ELEN

To systematically assess the capabilities of ELEN, we designed a series of experiments that address (i) structural sensitivity (including controlled geometric perturbations such as helical geometry scans, backbone and dihedral distortions, and systematic polar-to-hydrophobic mutations), (ii) sequence recognition and the contribution of LLM embeddings, and (iii) application-oriented benchmarking, which we do by comparing the performance of ELEN on challenging and diverse real-world protein modeling tasks to that of established methods. Full details of dataset construction, perturbation protocols, and evaluation metrics for each experiment are provided in the Supplementary Methods.

## 4. Results and Discussion

### 4.1 The ELEN loop dataset: coverage and diversity

We curated the ELEN loop dataset, a comprehensive and non-redundant resource comprising approximately 1 million protein loop pockets extracted from high-quality, diverse crystal structures. As summarized in Figure 2, the dataset exhibits high per-residue agreement between AF2 models and crystal structures (mean lDDT: 0.901, median: 0.938; mean CAD-score: 0.835, median: 0.87; mean RMSD: 0.161, median: 0.831). The length of the loops ranges from 2 to 10 residues, where shorter loops are more frequent. The amino acid composition reflects established loop-forming preferences. Importantly, each “loop pocket” includes both the loop and its surrounding structural context (Figure 2D, light magenta), which ensures that relevant contextual features are available for learning and predicting loop quality. The dataset is divided into training (938,000 samples), validation (186,000 samples), and testing (18,500 samples) sets, ensuring rigorous benchmarking. The ELEN loop dataset is available in both .pdb and LMDB formats at https://doi.org/10.5281/zenodo.15669210. Due to its large size (∼200 GB), the Saprot embeddings file is not publicly hosted but can be made available upon request to the corresponding author.

### 4.2 Performance evaluation on the CAMEO dataset

To comprehensively benchmark our ELEN models against other state-of-the-art MQA methods, we utilized a test set from the MQE category of the CAMEO online server. Performance was assessed on both loop residues and all residues (see Table 2), at both local (per-residue) and global (per-loop) levels. We report Spearman and Pearson correlation coefficients to assess ranking capability and accuracy of absolute quality predictions, respectively, alongside the Area Under the ROC Curve (AUC) for binary classification accuracy. Additionally, at the global (per-loop) level, we provide the Top-1 loss, defined as the average quality score difference between the predicted and actual best-performing model per target.

**Table 2.** Comparison of local and global MQA performance for ELEN, its ablation variants, and state-of-the-art predictors on the CAMEO MQE benchmark. Metrics are reported for both loop residues and all residues, and include Spearman’s (ρ), Pearson correlation coefficient (r), Area Under the ROC Curve (AUC), and Top-1 loss (difference between predicted and actual best-performing models per target). Results are shown separately for local (per-residue) and global (per-loop) evaluations. ELEN-GeomOnly denotes a geometry-only variant; ELEN-NoSeq excludes deep sequence embeddings. The highest values for each metric are highlighted in bold.

| Method | Loop residues |  |  |  |  |  |  | All residues |  |  |  |  |  |  |
| --- | --- | --- | --- | --- | --- | --- | --- | --- | --- | --- | --- | --- | --- | --- |
|  | Local MQA |  |  | Global MQA |  |  |  | Local MQA |  |  | Global MQA |  |  |  |
| | $\rho$ | $r$ | AUC | $\rho$ | $r$ | AUC | Top 1 | $\rho$ | $r$ | AUC | $\rho$ | $r$ | AUC | Top 1 |
| Baseline | 0.376 | 0.426 | 0.692 | 0.398 | 0.783 | 0.606 | 0.696 | 0.354 | 0.390 | 0.680 | 0.449 | 0.448 | 0.732 | 0.598 |
| ELEN | **0.657** | 0.717 | **0.818** | 0.744 | 0.842 | 0.939 | 0.617 | **0.688** | 0.737 | **0.839** | **0.749** | 0.829 | 0.840 | **0.468** |
| ELEN-NoSeq | 0.583 | 0.613 | 0.780 | **0.769** | 0.807 | **0.955** | 0.632 | 0.621 | 0.641 | 0.804 | 0.636 | 0.738 | 0.778 | 0.495 |
| ELEN-GeomOnly | 0.525 | 0.498 | 0.752 | 0.693 | **0.944** | 0.909 | 0.641 | 0.537 | 0.517 | 0.760 | 0.675 | 0.742 | 0.800 | 0.547 |
| ModFOLD7_IDDT | 0.382 | 0.487 | 0.684 | 0.391 | 0.545 | 0.864 | 0.641 | 0.421 | 0.566 | 0.711 | 0.430 | 0.668 | 0.675 | 0.503 |
| ModFOLD8 | 0.385 | 0.484 | 0.684 | 0.355 | 0.533 | 0.864 | 0.640 | 0.423 | 0.562 | 0.711 | 0.433 | 0.671 | 0.677 | 0.506 |
| ModFOLD9 | 0.628 | **0.727** | 0.812 | 0.700 | 0.895 | **0.955** | 0.609 | 0.639 | **0.761** | 0.824 | 0.748 | **0.837** | **0.858** | 0.480 |
| ProQ3 | 0.471 | 0.316 | 0.732 | 0.418 | 0.363 | 0.894 | 0.644 | 0.503 | 0.355 | 0.751 | 0.593 | 0.581 | 0.769 | 0.489 |
| ProQ3D | 0.412 | 0.420 | 0.700 | 0.564 | 0.748 | 0.879 | 0.664 | 0.444 | 0.379 | 0.716 | 0.535 | 0.620 | 0.742 | 0.519 |
| ProQ3D_LDDT | 0.474 | 0.591 | 0.737 | 0.769 | 0.876 | 0.879 | 0.628 | 0.504 | 0.633 | 0.752 | 0.564 | 0.774 | 0.761 | 0.500 |
| QMEANDisCo_3 | 0.485 | 0.584 | 0.741 | 0.630 | 0.657 | 0.879 | **0.606** | 0.517 | 0.636 | 0.767 | 0.532 | 0.745 | 0.732 | 0.503 |
| QMEAN_3 | 0.530 | 0.612 | 0.764 | 0.615 | 0.889 | 0.909 | 0.625 | 0.530 | 0.628 | 0.770 | 0.627 | 0.785 | 0.794 | 0.502 |
| VoroMQA_sw5 | 0.341 | 0.401 | 0.667 | 0.262 | 0.288 | 0.773 | 0.683 | 0.374 | 0.452 | 0.683 | 0.508 | 0.675 | 0.736 | 0.548 |
| VoroMQA_v2 | 0.490 | 0.563 | 0.739 | 0.537 | 0.813 | 0.788 | 0.664 | 0.546 | 0.555 | 0.776 | 0.636 | 0.666 | 0.792 | 0.556 |

For loop residues, ELEN achieved the highest Spearman rank correlation (0.657), which demonstrates its superior ability in correctly ranking residues by predicted quality. In contrast, ModFOLD9 attained the highest Pearson correlation (0.727), indicating slightly better linear predictions of absolute quality scores. At the global (per-loop) level, in terms of the Top-1 loss, ELEN was competitive, closely matching ModFOLD9 and QMEANDisCo_3, with QMEANDisCo_3 performing marginally better. Considering the other global metrics, vanilla ELEN model showed strong overall performance. However, other ELEN variants (see Table 1 for details) without sequence embeddings (ELEN-NoSeq) or with purely geometric features (ELEN-GeomOnly) exhibited unexpectedly higher performance on certain metrics. Specifically, ELEN-NoSeq achieved the highest global Spearman correlation (0.769) and AUC (0.955), while ELEN-GeomOnly recorded the highest global Pearson correlation (0.944). These findings suggest that at the global (per-loop) level, geometric features alone (ELEN-GeomOnly) or in combination with simplified sequence representations without explicit embeddings (ELEN-NoSeq) can yield unexpectedly robust predictive performance. This shows that detailed sequence embeddings provided by SaProt are less critical for capturing overall loop quality, potentially due to the dominance of geometric and physicochemical properties, such as Rosetta energies, hydrogen bonding, and solvent accessibility, in determining global structural quality.

These results collectively highlight ELEN’s consistent and competitive performance in protein MQA at both local and global levels, with particular strengths in correctly ranking residue quality. Importantly, unlike ModFOLD9 — a consensus method trained on the output of multiple other state-of-the-art MQA predictors — ELEN achieves strong performance without relying on any external model predictions, which underscores its generalizability and standalone effectiveness. The differing behaviors of ELEN variants at the global level between loop-specific and all-residue analyses provide valuable insights into the distinct predictive utilities of geometric and sequence-derived features in protein quality assessment at both local and global levels, with particular strengths in correctly ranking residue quality. The differing behaviors of ELEN variants at the global level between loop-specific and all-residue analyses provide valuable insights into the distinct predictive utilities of geometric and sequence-derived features in protein quality assessment.

### 4.3 Feature selection and ablation analysis

To determine the most informative input features, we systematically evaluated a wide range of per-residue and per-atom descriptors — including continuous (Rosetta energies, SASA, SAP-score), categorical (sequence, secondary structure), and LLM-derived sequence embeddings (e.g. SaProt_650M_PDB). The addition of continuous descriptors at both atom and residue levels, particularly Rosetta energies and SASA, consistently improved performance (mean Pearson’s r up to 0.37). Notably, sequence embeddings from SaProt provided the greatest enhancement, yielding the highest observed correlation (r = 0.44). Several per-atom features (e.g., number of hydrogen bonds per atom, atomic B-factors, and charges) were also tested, but were ultimately excluded due to their computational complexity and limited additional predictive benefit relative to per-residue descriptors. Full ablation results and detailed performance metrics are provided in Supplementary Table S2 and Supplementary Figure S1. Collectively, these findings guided the final ELEN model design, which integrates both geometric and physicochemical features as well as sequence embeddings for optimal loop quality prediction. The complete set of features included in the ELEN model and its ablation variants is summarized in Table 1.

### 4.4 Structural and geometric awareness of ELEN

To assess ELEN’s ability to recognize structural quality, we performed three analyses focused on helical geometry, backbone distortions, and hydrogen bond network disruption (Figure 3). For comparison, the performance of the sequence-agnostic (ELEN-NoSeq) and geometry-only (ELEN-GeomOnly) variants is shown in Figures S2–S4 and Tables S3–S4 in the Supplementary. Importantly, ELEN scores are not calibrated across models and, thus, our analysis focuses on relative score changes (sensitivity in this context) within each variant.

**Figure 3.**
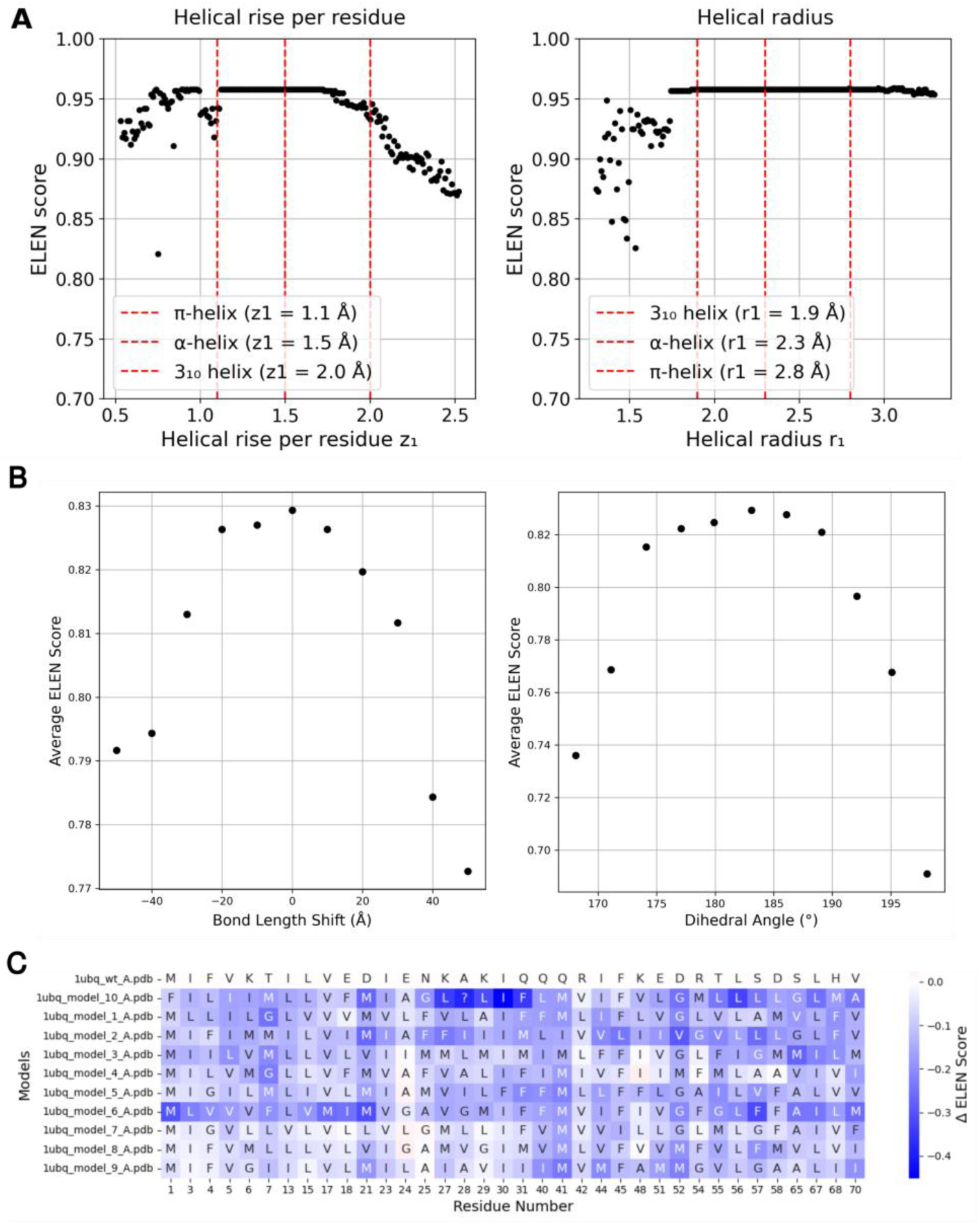
Integrated assessment of the geometric and structural sensitivity of ELEN. (A) Helical geometry selectivity: The mean per-chain ELEN score is plotted as a function of helical rise per residue (z_1_, left) and helical radius (r_1_, right) for a series of idealized 25-residue helices generated with systematically varied helical parameters. Vertical red dashed lines indicate canonical values for π-helix, α-helix, and 3_10_-helix geometries, highlighting each model’s sensitivity and selectivity for native-like helical structure. (B) Backbone perturbation sensitivity: Average ELEN scores for residues 61–63 of ubiquitin (PDB ID: 1UBQ) are plotted as a function of backbone bond length shift (left) and peptide bond dihedral angle (ω) (right) for the full ELEN model. Decreasing ELEN scores indicate loss of structural plausibility as deviations from native geometry increase. (C) Impact of polar-to-hydrophobic mutations: Heatmaps show changes in per-residue ELEN scores (ΔELEN) relative to the wild-type structure for ubiquitin following systematic replacement of polar residues (forming at least two hydrogen bonds) with hydrophobic amino acids. Blue indicates a reduction in predicted structural quality, while red denotes a relative improvement, reflecting each model’s sensitivity to disruption of hydrogen bonding networks.

#### 4.4.1 Helical geometry preferences

We evaluated ELEN’s sensitivity to secondary structure geometry by scoring a series of idealized helices in which the helical rise per residue (z_1_) and helical radius (r_1_) were systematically varied. For both parameters, ELEN assigned maximal scores at values closely matching those of canonical α-, 3_10_-, and π-helices (Figure 3A), with scores dropping sharply as geometries deviated from these native-like values. This demonstrates that ELEN is finely tuned to recognize and prefer natural helical conformations. The NoSeq and GeomOnly variants (Supplementary Figures S2B and S2C) displayed broadly similar trends, which confirms that geometric features alone confer substantial discriminatory power. However, the ELEN-GeomOnly model showed increased score variability, with less pronounced selectivity, particularly for the helical radius. Notably, ELEN-GeomOnly often assigned relatively high scores to helices whose radii are far outside the range of radii that is typically observed in natural proteins. These findings suggest that while geometric information is essential, the absence of sequence context reduces the model’s ability to distinguish biologically plausible from merely physically possible geometries. Consequently, ELEN-GeomOnly is more permissive and prone to scoring non-natural conformations highly, which reflects a loss of evolutionary and physicochemical discrimination. Quantitative comparisons of mean ELEN scores across all model variants and helical parameter sets are provided in Supplementary Table S3. Overall, these results underscore the complementary roles of geometric and sequence features in robust secondary structure assessment.

#### 4.4.2 Sensitivity of ELEN to backbone and dihedral distortions

To assess the sensitivity of ELEN and its variants to local structural distortions, we systematically perturbed backbone bond lengths and peptide bond dihedral angles in ubiquitin (Supplementary Figure S3D). All ELEN models responded to deviations from native geometry, but with notable differences in sensitivity (Supplementary Figures S3A–C). The full ELEN model exhibited a pronounced decrease in average per-residue score as bond lengths and dihedral angles deviated from their canonical values (Figure 3B), which indicates strong discrimination against unphysical backbone conformations. This sensitivity was especially marked for extreme distortions, which underscores ELEN’s capacity to detect highly improbable geometries. The ELEN-NoSeq variant showed a qualitatively similar but somewhat attenuated response, which suggests that sequence information enhances the, but is not strictly required for, recognition of backbone anomalies. In contrast, the ELEN-GeomOnly model exhibited much less variation in scores across the perturbation range (Supplementary Figure S3C), which indicates a limited ability to identify structural anomalies when both sequence and comprehensive structural context are absent. Overall, these results demonstrate that both geometric and sequence features contribute to ELEN’s ability to robustly detect deviations from canonical backbone geometry, with sequence information conferring additional sensitivity to subtle structural perturbations.

#### 4.4.3 Impact of polar-to-hydrophobic mutations on ELEN scores

To assess the influence of hydrogen bonds on ELEN predictions, we systematically replaced polar, hydrogen bond-forming residues in ubiquitin (PDB ID: 1UBQ) and a *de novo* four-helix bundle (PDB ID: 4UOS) with hydrophobic amino acids, using thresholds based on the number of hydrogen bonds formed (Supplementary Table S4). Mutations were performed for residues forming at least one, two, or three hydrogen bonds in the wild-type structures, and the resulting models were evaluated using the full ELEN model, as well as the sequence-agnostic (ELEN-NoSeq) and geometry-only (ELEN-GeomOnly) variants. Across all cases, mutations consistently reduced the predicted per-residue quality as measured by the full ELEN model (Figure 4C). In ubiquitin, the largest decrease in ELEN score was observed for mutations at the lowest threshold (≥1 hydrogen bond, mean ΔELEN = -0.2695 ± 0.0960), with the effect diminishing as the threshold increased (≥2: -0.1214 ± 0.0730; ≥3: -0.0545 ± 0.0230). A similar trend was seen for the four-helix bundle (≥1: -0.1765 ± 0.0900; ≥2: -0.1368 ± 0.0762), while mutations of residues involved in three or more hydrogen bonds had minimal impact. The smaller effect in the four-helix bundle likely reflects its inherently more robust and mutation-tolerant design. Per-residue analysis at the ≥2 hydrogen bond threshold (Figure 3C) revealed that the full ELEN model detected a pronounced and widespread reduction in predicted quality, while the ELEN-NoSeq and ELEN-GeomOnly variants (Supplementary Figures S4B, S4C) showed more localized and less pronounced changes, occasionally even predicting improvements. These results highlight the enhanced sensitivity of the full ELEN model and underscore the value of integrating sequence-specific and physicochemical features for detecting the structural consequences of hydrogen bond disruption.

### 4.5 Assessing ELEN in AF2 failure cases and advanced design scenarios

#### 4.5.1 Benchmarking ELEN on challenging AF2 targets

To investigate ELEN’s effectiveness in identifying structural inaccuracies in protein models, we benchmarked it on twelve structurally diverse protein targets that were previously identified as challenging to predict for AF2. These targets, curated from [37], encompass various structural classes, including peptides, membrane proteins, intrinsically disordered proteins, multi domain complexes, and oligomers. Each AF2 model was evaluated using ELEN, and per-residue quality scores were visualized by color-coding on the predicted structures, which were then superimposed onto their experimental counterparts (Figure 5). Visual inspection revealed that ELEN effectively highlighted structurally inaccurate regions, such as locally misfolded regions, loop misplacements, and intrinsically disordered segments. Quantitative evaluation (Supplementary Table S5) confirmed these observations: ELEN identified the main structural flaw (defined by a score cutoff of 0.8) in at least seven out of twelve cases, whereas AF2’s intrinsic confidence metric, plDDT (cutoff 70), achieved this for only four targets. Specifically, ELEN consistently achieved higher overlap and Jaccard indices (a measure for the similiarity of two sets) compared to plDDT in cases that involved pronounced local misfolding and disorder. Notably, for targets characterized by significant misfolding or intrinsic disorder (e.g., cases 2KQP, 6TUB, and 1S4T), ELEN strongly outperformed plDDT, which emphasizes its utility for accurately identifying flexible or disordered structural elements. However, both ELEN and plDDT showed limited specificity for large-scale domain orientation errors (Figure 5D, 5E, 5K), as these metrics inherently focus on local, per-residue accuracy rather than global domain arrangements. These findings underscore a fundamental limitation of residue-based scoring methods and highlight the need for complementary global metrics (e.g., TM-score or domain-level RMSD) to comprehensively assess protein structure quality. Overall, ELEN’s enhanced sensitivity in detecting localized inaccuracies complements existing metrics such as plDDT, making it particularly valuable for targeted refinement and validation of protein structure predictions.

**Figure 5.**
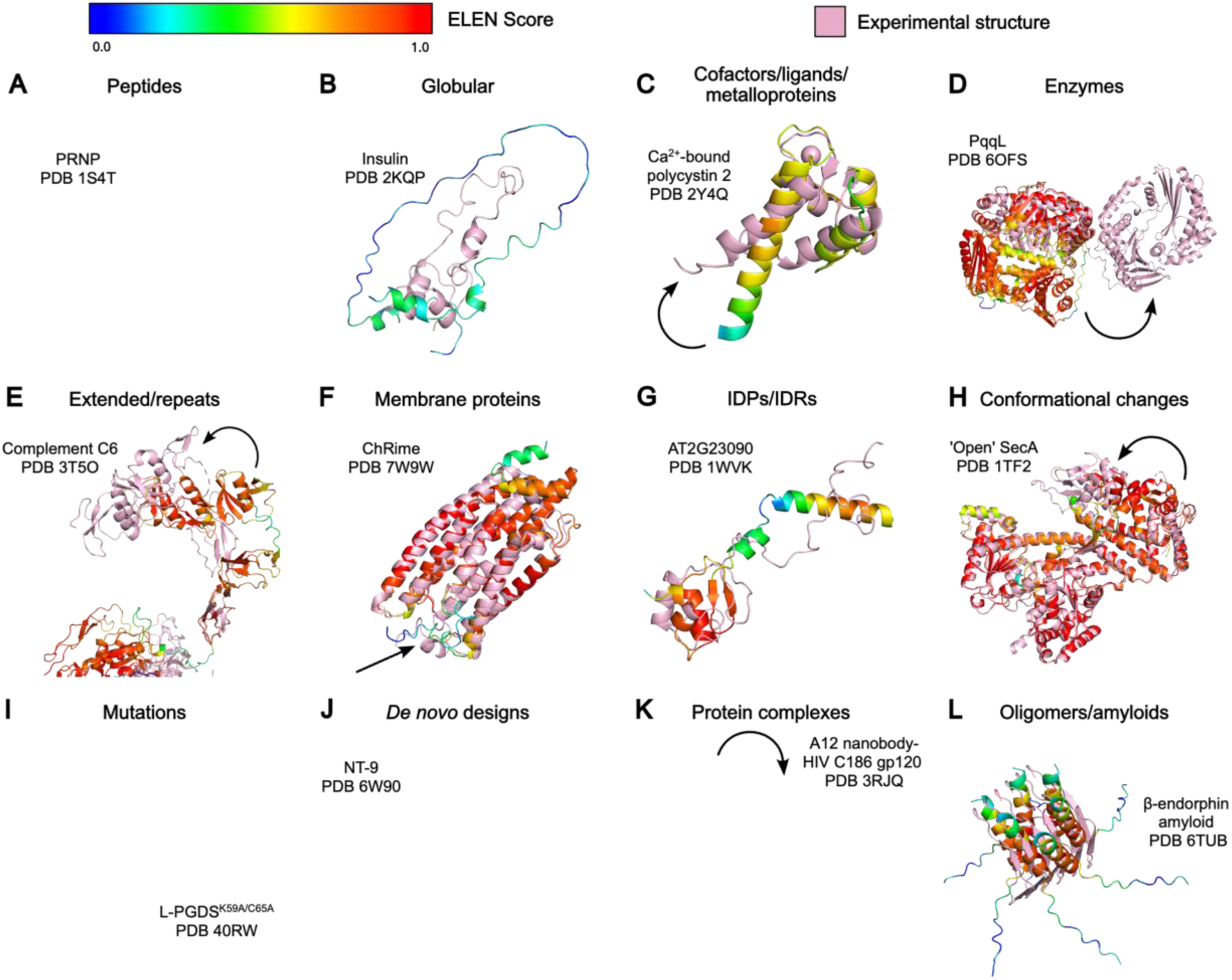
Visualization of ELEN predicted local model quality for AF2 predictions that deviate from the experimental structure. AF2 models for twelve proteins, selected from the benchmark dataset presented in [37], are shown superimposed on their respective experimental structures (pink). AF2 models are colored according to ELEN per-residue quality scores, from low (blue) to high (red) as indicated by the color bar. Peptide (A), insulin (B), polycystin 2 (C), PqqL (D), complement component C6 (E), ChRmine (F), AT2G23090 (G), SecA (H), lipocalin-type PGDS (L-PGDS)K59A/C65A (I), NT-9 (J), A12 nanobody–HIV C186 gp120 complex (K), and β-endorphin amyloid fibril (L). Black arrows highlight regions where the AF2 model deviates from the crystal structure.

#### 4.5.2 Assessing ELEN for the detection of enhanced loop design

To determine whether ELEN can effectively distinguish between low- and high-quality designed loops, we employed ELEN to optimize loop variants of the human duodenal cytochrome b (Dcytb) protein (see Supplementary). Our evaluation focused on the important loop between residues 180–190, and the remainder of the structure, using three complementary metrics: the ELEN score, AlphaFold plDDT, and all-atom RMSD to the native crystal structure (PDB ID: 5ZLG). Summary statistics for all regions and design sets are provided in Supplementary Table S6. ELEN guided redesigned loops consistently demonstrated higher plDDT values than the original designs. Specifically, in the loop region, it was possible to increase the average ELEN score from 0.812 (original) to 0.839 (redesign), and the mean plDDT from 70.3 to 82.4. Interestingly, while the redesigned loops exhibit greater predicted local quality and model confidence, they adopt alternative conformations that are different from the wild-type loop (RMSD to wild type 5.64Å). This pattern persisted when assessing the rest of the protein and the entire structure: Both ELEN and plDDT scores were significantly higher in redesigned models, while RMSD differences outside the loop were comparatively modest. These findings collectively suggest that the new loop designs achieve improved computational quality and confidence yet diverge structurally from the starting scaffold. The distributions of ELEN scores, plDDT, and RMSD for the loop region are visualized in Figure 6B, where violin plots with overlaid boxplots reveal a marked shift toward higher ELEN and plDDT values in redesigned loops, alongside a broader and elevated RMSD distribution. Figure 6A presents representative structure overlays of the lowest- and highest-scoring loop designs (colored by ELEN score) relative to the native reference (pink). As a negative control, the native crystal structure was also evaluated using ELEN (Figure 6A, right). In summary, these results highlight the sensitivity of ELEN to loop design quality and confirm that redesigned loops are predicted to possess superior quality and confidence.

**Figure 6.**
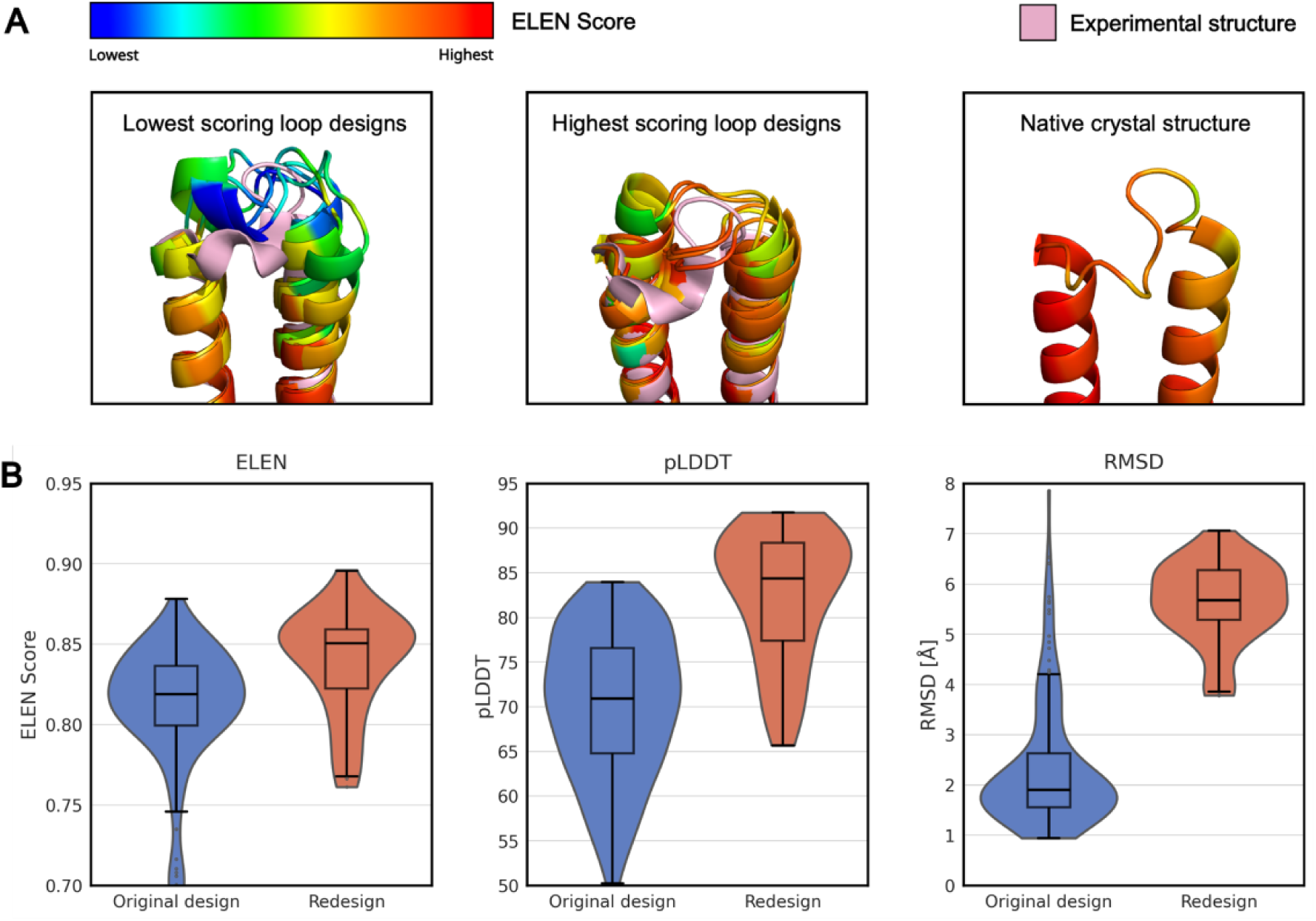
Structural and quantitative comparison of the original and redesigned loop region (residues 180-190) in human Dcytb. (A), Representative structures of the five worst-performing (left) and five best-performing (middle) loop designs colored with respect to the ELEN score. The native crystal structure is shown for reference (right). (B), Violin plots with overlaid box plots compare ELEN score, plDDT, and loop RMSD distributions between original (blue) and redesigned (orange) loops for residues 180–190. Horizontal lines within each violin represent the median and interquartile range. Distributions reflect all models analyzed (original: n = 161; redesign: n = 52).

#### 4.5.3 ELEN detects disordered and variable regions in *de novo* designed enzymes

We evaluated the performance of ELEN on a dataset of *de novo* designed Kemp eliminases to determine its ability to predict model accuracy, identify structurally variable regions, and specifically detect disordered segments that lack defined electron density in experimentally determined crystal structures. For one representative Kemp eliminase, an experimentally determined crystal structure was available (PDB ID: 9HVB), which is known to lack clear electron density in three distinct regions, indicating local disorder. Figure 7 (left) shows a structural overlay of the experimentally determined crystal structure (purple) and a representative computational model, with the latter colored according to per-residue ELEN scores. Importantly, ELEN assigns lower scores (yellow) to the regions that are missing in the crystal structure, which correspond precisely to the experimentally unresolved segments (annotated as Disordered regions 1–3). This agreement demonstrates that ELEN is sensitive to poor local backbone-sidechain pairs, and can highlight segments where structural models are intrinsically unreliable due to physical flexibility or bad model quality. Supplementary Figure S5 presents a direct, residue-by-residue comparison between ELEN scores across the ensemble of computational models, the crystal structure scored by ELEN, and the experimental B-factors. Regions of the crystal structure with low B-factors — indicative of well-ordered, rigid residues — consistently correspond to stretches of low ELEN scores in the models, which demonstrates both the accuracy of the design protocol in stabilizing the protein core and ELEN’s sensitivity to structural reliability. In contrast, regions with elevated B-factors, typically associated with loops and solvent-exposed segments, manifest as vertical bands of high ELEN scores across models, which reflects on increased local uncertainty and flexibility. Importantly, residues that lack electron density in the crystal structure (annotated as ‘X’) are systematically assigned higher ELEN scores across the ensemble, which underscores the capacity of ELEN to detect sub-optimal parts of the model, which may correspond disordered regions. The robust co-localization of high ELEN scores with regions of missing density highlights model-intrinsic flaws that may impair catalytic performance or protein production. This provides actionable insights for further optimization. Quantitative analysis, summarized in Supplementary Table S7, further underscores these findings. The mean ELEN scores for residues within the disordered regions are significantly higher than for those resolved in the crystal structure, with differences ranging from ∼0.024 to 0.038 across the three disordered segments. In contrast, the ELEN score of the crystal structure was uniformly low and showed low variance, as expected for a well-defined experimental structure. Importantly, the close agreement between mean ELEN scores and experimental B-factors in resolved regions supports ELEN’s utility as a surrogate measure of local order and model reliability. This significant quantitative agreement highlights ELEN’s value for identifying sites of disorder and guiding targeted refinement in protein design. Taken together, these results establish ELEN as a powerful tool for pinpointing both resolved and problematic regions in protein models. The method not only mirrors experimental disorder but also exposes sequence segments where further stabilization or redesign may be warranted. By enabling direct, per-residue comparison between predicted model uncertainty and experimental data, ELEN can guide targeted stability engineering—such as via PROSS or FuncLib protocols—to optimize both the global fold and catalytic function of designed enzymes. Furthermore, this approach enables the identification of synergistic mutational effects, both within and outside active sites, facilitating rational improvement of protein designs. Conversely, discrepancies between ELEN and experimental profiles highlight sites for further refinement.

To further demonstrate ELEN’s capability to detect deviations from experimentally determined structures, we analyzed a set of de novo retro-aldolases designs (RAD) for which both a high-resolution crystal structure and an AF3 model were available. In this case, a distinct loop region displays a significant deviation between the AF3 model and the crystal structure, with the computational model adopting a conformation markedly different from experiment (Figure 7, right). Strikingly, ELEN assigns lower per-residue scores to this segment and accurately flags it as an area of reduced model reliability. Notably, the crystallographic B-factors for this loop do not indicate increased flexibility, which suggests that the structural discrepancy arises from modeling inaccuracies rather than intrinsic disorder or mobility.

**Figure 7.**
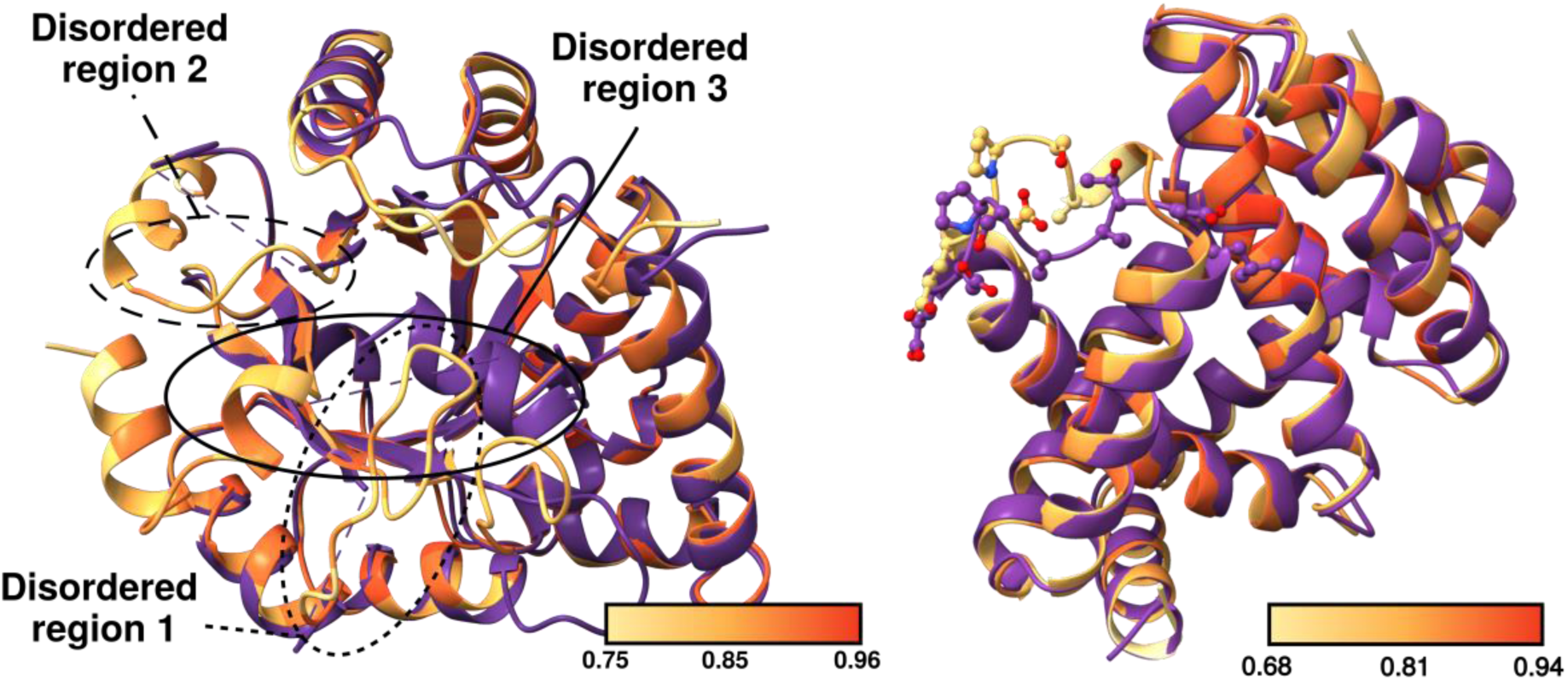
ELEN detects local deviations between computational models and experimental structures. Structural overlays of the crystal structure (purple) and computational models (orange–yellow, colored by per–residue ELEN scores; higher scores indicate higher predicted local quality) are shown for two designed proteins. Left: A de novo designed Kemp eliminase, where regions absent from the crystal structure due to missing electron density (annotated as regions 1–3) correspond to stretches of low model quality predicted by ELEN, consistent with experimental disorder. Right: A RAD protein, in which a loop region exhibiting a pronounced deviation between the AlphaFold3 model and the crystal structure is flagged by elevated ELEN scores, highlighting a localized modeling inaccuracy.

## 5 Interpreting ELEN’s internal representations with PCA

To evaluate which structural and physicochemical properties are encoded by the ELEN model, we performed principal component analysis (PCA) on the penultimate layer activations for all residues across the CAMEO one-month dataset and colored the resulting projections according to various pre-computed residue-level features. To further investigate how different input features influence these learned representations, we compared the full ELEN model with the geometry-only variant, ELEN-GeomOnly (Supplementary Figure S6). In the full ELEN model, side-chain identity and ABEGO type (Supplementary Figure 6A, 6B) exhibit partial clustering, which indicates that the model encodes sequence and backbone information robustly. Continuous features such as per-residue SASA (Supplementary Figure 6C), number of hydrogen bonds (Supplementary Figure 6D), backbone dihedral angles (Supplementary Figure S6E, S6F), and Rosetta energy (Supplementary Figure S6G) exhibit clear gradients or localized regions in the PCA space, which suggests that these physicochemical features are well represented in the learned embeddings. As a negative control, residue index was included (Supplementary Figure S6H). Consistent with our expectations, the residue index does not carry biologically relevant structural information and hence *–* in contrast to the true structural features *–* did not show pronounced clustering or spatial gradients in the PCA projection.

The ELEN-GeomOnly model, which was trained without additional physicochemical features and SaProt sequence embeddings, displays less pronounced color separation and weaker gradients for the majority of features. In particular, the encoding of side-chain identity and ABEGO pattern was markedly reduced compared to the full model. This reduction indicates that the model’s ability to represent and disentangle physicochemical properties in its learned representations is strongly influenced by the inclusion of these features during training. Overall, these results demonstrate that ELEN’s learned representations capture a combination of geometric, energetic, and sequence-derived features at the residue level, supporting the model’s capacity to integrate diverse structural signals relevant for protein quality assessment.

## 6 Conclusion

We introduce ELEN, a deep learning-based MQA method specifically designed for precise, per-residue quality assessment of protein loops at atomic resolution. By simultaneously predicting lDDT, CAD-score, and RMSD, and integrating physicochemical features and sequence embeddings derived from LLMs, ELEN achieves state-of-the-art performance, surpassing existing single-model MQA methods on the challenging CAMEO benchmark. We also show ELEN’s practical applicability by performing detailed analysis, including identification of flexible or disordered regions and assessment of structural effects of single-residue variants in redesigned enzymes. Our work is underpinned by the ELEN loop dataset—a large, diverse, and high-quality resource of over one million loop pockets from experimental structures. Both the dataset and code are freely available to support further advances in protein modeling. Ablation analyses revealed that ELEN variants leveraging only geometric or physicochemical features often perform surprisingly well, underlining the critical contribution of local geometry and physicochemical properties to model quality.

ELEN can detect subtle local inaccuracies—including hydrogen bond disruptions and structural irregularities—highlighting poorly designed regions, validating improved loop redesigns, and identifying disordered segments lacking electron density. These capabilities underscore its practical value for protein engineering and design.

Although the current version of ELEN is trained exclusively on static, single-chain crystal structures and does not explicitly model ligands or structural flexibility, our findings lay a robust groundwork for future enhancements. Expanding training to include ligand-containing AF3 models, data from molecular dynamics simulations, and multi-chain assemblies will further extend ELEN’s applicability. Overall, ELEN represents a robust, flexible, and interpretable framework for assessing protein loop quality, offering substantial advancements for computational protein design.

## 7 Data Availability

### Materials & Correspondence

Supplementary Materials are available for this paper. The training dataset generated and analyzed in this study is publicly available via Zenodo at https://doi.org/10.5281/zenodo.15669210. Due to file size, sequence embedding files (SaProt) are available upon reasonable request to the corresponding author. The code for ELEN model inference is accessible at https://github.com/FlorianWieser1/ELEN_inference and is distributed under an open-source license. Portions of this manuscript were drafted and revised with the assistance of OpenAI’s ChatGPT language model, as required by the publisher’s guidelines. Correspondence and requests for materials should be addressed to G.O.

## 8 Acknowledgments

## Acknowledgments

We thank Andreas Winkler, Johannes Peterlechner and Allon Hochbaum for discussions about the approach and interpretability of the results. We also thank Shlomo Yakir Hoch and Sarel Fleishman for providing us early access to the model and crystal structure of Kemp Eliminase KE (Des27.7) for ELEN analysis. Additionally, we acknowledge the Graz University of Technology high-performance computing facility for access to compute. For open access, the authors have applied a CC BY public copyright license to any Author Accepted Manuscript version arising from this submission.

## Author contributions

Conceptualization: F.W., G.O.; Methodology: F.W., M.Z., T.P., G.O.; Software: F.W., M.Z.; Validation: F.W., S.K.; Formal analysis: F.W., S.K.; Investigation: F.W., S.K., M.Z., G.O.; Resources: T.P., G.O.; Data Curation: F.W., S.K.; Writing: F.W., G.O.; Visualization: F.W., S.K.; Supervision: T.P., G.O.; Project administration: G.O.; Funding acquisition: G.O.

## References

[1] S. Kwon, J. Won, A. Kryshtafovych, and C. Seok, “Assessment of protein model structure accuracy estimation in CASP14: Old and new challenges,” Proteins, vol. 89, no. 12, pp. 1940–1948, 2021, doi: 10.1002/prot.26192.

[2] J. Jumper et al., “Highly accurate protein structure prediction with AlphaFold,” Nature, vol. 596, no. 7873, pp. 583–589, Aug. 2021, doi: 10.1038/s41586-021-03819-2.

[3] J. Abramson et al., “Accurate structure prediction of biomolecular interactions with AlphaFold 3,” Nature, vol. 630, no. 8016, pp. 493–500, June 2024, doi: 10.1038/s41586-024-07487-w.

[4] M. Baek et al., “Accurate prediction of protein structures and interactions using a three-track neural network,” Science, vol. 373, no. 6557, pp. 871–876, Aug. 2021, doi: 10.1126/science.abj8754.

[5] Z. Lin et al., “Evolutionary-scale prediction of atomic level protein structure with a language model,” July 21, 2022. doi: 10.1101/2022.07.20.500902.

[6] T. Hayes et al., “Simulating 500 million years of evolution with a language model,” Science, vol. 387, no. 6736, pp. 850–858, Feb. 2025, doi: 10.1126/science.ads0018.

[7] C. Chen, X. Chen, A. Morehead, T. Wu, and J. Cheng, “3D-equivariant graph neural networks for protein model quality assessment,” Bioinformatics, vol. 39, no. 1, p. btad030, Jan. 2023, doi: 10.1093/bioinformatics/btad030.

[8] A. O. Stevens and Y. He, “Benchmarking the Accuracy of AlphaFold 2 in Loop Structure Prediction,” Biomolecules, vol. 12, no. 7, p. 985, July 2022, doi: 10.3390/biom12070985.

[9] T. C. Terwilliger et al., “AlphaFold predictions are valuable hypotheses and accelerate but do not replace experimental structure determination,” Nat Methods, vol. 21, no. 1, pp. 110–116, 2024, doi: 10.1038/s41592-023-02087-4.

[10] D. Liu, B. Zhang, J. Liu, H. Li, L. Song, and G. Zhang, “Assessing protein model quality based on deep graph coupled networks using protein language model,” Briefings in Bioinformatics, vol. 25, no. 1, p. bbad420, Nov. 2023, doi: 10.1093/bib/bbad420.

[11] L. J. McGuffin, F. M. F. Aldowsari, S. M. A. Alharbi, and R. Adiyaman, “ModFOLD8: accurate global and local quality estimates for 3D protein models,” Nucleic Acids Research, vol. 49, no. W1, pp. W425–W430, July 2021, doi: 10.1093/nar/gkab321.

[12] S.-S. Guo, J. Liu, X.-G. Zhou, and G.-J. Zhang, “DeepUMQA: ultrafast shape recognition-based protein model quality assessment using deep learning,” Bioinformatics, vol. 38, no. 7, pp. 1895–1903, Mar. 2022, doi: 10.1093/bioinformatics/btac056.

[13] K. Uziela, D. Menéndez Hurtado, N. Shu, B. Wallner, and A. Elofsson, “Improved protein model quality assessments by changing the target function,” Proteins, vol. 86, no. 6, pp. 654–663, 2018, doi: 10.1002/prot.25492.

[14] A. H. A. Maghrabi and L. J. McGuffin, “Estimating the Quality of 3D Protein Models Using the ModFOLD7 Server,” in Protein Structure Prediction, vol. 2165, D. Kihara, Ed., New York, NY: Springer US, 2020, pp. 69–81. doi: 10.1007/978-1-0716-0708-4_4.

[15] A. Ray, E. Lindahl, and B. Wallner, “Improved model quality assessment using ProQ2,” BMC Bioinformatics, vol. 13, no. 1, p. 224, 2012, doi: 10.1186/1471-2105-13-224.

[16] L. J. McGuffin and S. M. A. Alharbi, “ModFOLD9: A Web Server for Independent Estimates of 3D Protein Model Quality,” Journal of Molecular Biology, p. 168531, 2024, doi: 10.1016/j.jmb.2024.168531.

[17] S. Eismann, P. Suriana, B. Jing, R. J. L. Townshend, and R. O. Dror, “Protein model quality assessment using rotation-equivariant transformations on point clouds,” Proteins, vol. 91, no. 8, pp. 1089–1096, 2023, doi: 10.1002/prot.26494.

[18] T. S. Cohen and M. Welling, “Group Equivariant Convolutional Networks,” 2016, doi: 10.48550/ARXIV.1602.07576.

[19] S. Eismann, P. Suriana, B. Jing, R. J. L. Townshend, and R. O. Dror, “Protein model quality assessment using rotation-equivariant, hierarchical neural networks,” 2020, arXiv. doi: 10.48550/ARXIV.2011.13557.

[20] V. Mariani, M. Biasini, A. Barbato, and T. Schwede, “lDDT: a local superposition-free score for comparing protein structures and models using distance difference tests,” Bioinformatics, vol. 29, no. 21, pp. 2722–2728, Nov. 2013, doi: 10.1093/bioinformatics/btt473.

[21] K. Olechnovič, E. Kulberkytė, and Č. Venclovas, “CAD-score: A new contact area difference-based function for evaluation of protein structural models,” Proteins, vol. 81, no. 1, pp. 149–162, 2013, doi: 10.1002/prot.24172.

[22] K. Olechnovič, B. Monastyrskyy, A. Kryshtafovych, and Č. Venclovas, “Comparative analysis of methods for evaluation of protein models against native structures,” Bioinformatics, vol. 35, no. 6, pp. 937–944, Mar. 2019, doi: 10.1093/bioinformatics/bty760.

[23] J. Haas et al., “Continuous Automated Model EvaluatiOn (CAMEO) complementing the critical assessment of structure prediction in CASP12,” Proteins, vol. 86, no. S1, pp. 387–398, 2018, doi: 10.1002/prot.25431.

[24] G. Wang and R. L. Dunbrack, “PISCES: a protein sequence culling server,” Bioinformatics, vol. 19, no. 12, pp. 1589–1591, Aug. 2003, doi: 10.1093/bioinformatics/btg224.

[25] H. M. Berman, “The Protein Data Bank,” Nucleic Acids Research, vol. 28, no. 1, pp. 235–242, Jan. 2000, doi: 10.1093/nar/28.1.235.

[26] M. Mirdita, K. Schütze, Y. Moriwaki, L. Heo, S. Ovchinnikov, and M. Steinegger, “ColabFold: making protein folding accessible to all,” Nat Methods, vol. 19, no. 6, pp. 679–682, 2022, doi: 10.1038/s41592-022-01488-1.

[27] W. Kabsch and C. Sander, “Dictionary of protein secondary structure: Pattern recognition of hydrogen-bonded and geometrical features,” Biopolymers, vol. 22, no. 12, pp. 2577–2637, 1983, doi: 10.1002/bip.360221211.

[28] S. J. Fleishman et al., “RosettaScripts: A Scripting Language Interface to the Rosetta Macromolecular Modeling Suite,” PLoS ONE, vol. 6, no. 6, p. e20161, June 2011, doi: 10.1371/journal.pone.0020161.

[29] K. Uziela, N. Shu, B. Wallner, and A. Elofsson, “ProQ3: Improved model quality assessments using Rosetta energy terms,” Sci Rep, vol. 6, no. 1, p. 33509, Oct. 2016, doi: 10.1038/srep33509.

[30] K. Uziela, D. Menéndez Hurtado, N. Shu, B. Wallner, and A. Elofsson, “ProQ3D: improved model quality assessments using deep learning,” Bioinformatics, vol. 33, no. 10, pp. 1578–1580, May 2017, doi: 10.1093/bioinformatics/btw819.

[31] G. Studer, C. Rempfer, A. M. Waterhouse, R. Gumienny, J. Haas, and T. Schwede, “QMEANDisCo—distance constraints applied on model quality estimation,” Bioinformatics, vol. 36, no. 6, pp. 1765–1771, Mar. 2020, doi: 10.1093/bioinformatics/btz828.

[32] P. Benkert, S. C. E. Tosatto, and D. Schomburg, “QMEAN: A comprehensive scoring function for model quality assessment,” Proteins, vol. 71, no. 1, pp. 261–277, 2008, doi: 10.1002/prot.21715.

[33] “Proteins - 2017 - Olechnovič - VoroMQA Assessment of protein structure quality using interatomic contact areas.pdf.”

[34] J. Su, C. Han, Y. Zhou, J. Shan, X. Zhou, and F. Yuan, “SaProt: Protein Language Modeling with Structure-aware Vocabulary,” Oct. 02, 2023. doi: 10.1101/2023.10.01.560349.

[35] E. C. Meng et al., “UCSF ChimeraX: Tools for structure building and analysis,” Protein Science, vol. 32, no. 11, p. e4792, 2023, doi: 10.1002/pro.4792.

[36] J. Dauparas et al., “Robust deep learning–based protein sequence design using ProteinMPNN,” Science, vol. 378, no. 6615, pp. 49–56, Oct. 2022, doi: 10.1126/science.add2187.

[37] V. Agarwal and A. C. McShan, “The power and pitfalls of AlphaFold2 for structure prediction beyond rigid globular proteins,” Nat Chem Biol, vol. 20, no. 8, pp. 950–959, 2024, doi: 10.1038/s41589-024-01638-w.

[38] D. Listov et al., “High-efficiency Kemp eliminases by complete computational design,” Jan. 04, 2025. doi: 10.1101/2025.01.04.631280.

[39] W. Kabsch, “A solution for the best rotation to relate two sets of vectors,” Acta Cryst A, vol. 32, no. 5, pp. 922–923, Sept. 1976, doi: 10.1107/S0567739476001873.

[40] M. M. Bronstein, J. Bruna, T. Cohen, and P. Veličković, “Geometric Deep Learning: Grids, Groups, Graphs, Geodesics, and Gauges,” 2021, arXiv. doi: 10.48550/ARXIV.2104.13478.

[41] J. K. Leman et al., “Macromolecular modeling and design in Rosetta: recent methods and frameworks,” Nat Methods, vol. 17, no. 7, pp. 665–680, 2020, doi: 10.1038/s41592-020-0848-2.

[42] J. Kurniawan and T. Ishida, “Protein Model Quality Estimation Using Molecular Dynamics Simulation,” ACS Omega, vol. 7, no. 28, pp. 24274–24281, July 2022, doi: 10.1021/acsomega.2c01475.

[43] G. Derevyanko, S. Grudinin, Y. Bengio, and G. Lamoureux, “Deep convolutional networks for quality assessment of protein folds,” Bioinformatics, vol. 34, no. 23, pp. 4046–4053, Dec. 2018, doi: 10.1093/bioinformatics/bty494.

