## Supplementary Materials for "ELEN – Predicting Loop Quality in Protein Structure Models"

**Content:**

Supplementary Methods: p. 2- p. 4
Supplementary Figures: S1 – S6
Supplementary Tables: S1 – S7

### Supplementary Methods: Experimental evaluation of ELEN

*Helical geometry preferences*

To assess ELEN's ability to recognize physically realistic helical geometries, we designed 200 helices, each 25 amino acids long, using the MakeBundle mover of RosettaScripts [28]. Two key parameters were systematically varied: the helical radius ($r_1$) and the helical rise per residue ($z_1$). The helical radius was adjusted from 1.53 Å to 3.06 Å, while the rise per residue ranged from 0.75 Å to 2.33 Å in 0.01 Å increments. Helices were then scored by the ELEN model and its variants, and the resulting scores were plotted as a function of these parameters.

*Sensitivity of ELEN to backbone and dihedral distortions*

To evaluate the sensitivity of ELEN to deviations from physically plausible protein backbone geometries, we performed systematic perturbations of both bond lengths and dihedral angles on the model protein ubiquitin (PDB ID: 1UBQ). Structural distortions were introduced using UCSF ChimeraX [35]. For bond length perturbations, the covalent bonds N–Cα, Cα–C, and C–N of residue 62 were incrementally lengthened or shortened in 0.1 Å steps relative to their canonical values. For dihedral angle perturbations, the peptide bond dihedral angle (ω) between residues 62 and 63 was incrementally increased or decreased in 3° steps. After each modification, the perturbed structures were assessed using the ELEN model and its variants to quantify the model's sensitivity to progressively unphysical backbone conformations. ELEN scores were subsequently averaged over residues 61, 62, and 63, and plotted as a function of the modified backbone parameter.

*Impact of polar-to-hydrophobic mutations on ELEN scores*

To assess the effect of hydrogen bond participation on ELEN model predictions, we performed systematic polar-to-hydrophobic mutations in two protein systems: the natural protein ubiquitin (PDB ID: 1UBQ) and a *de novo* designed four-helix bundle (PDB ID: 4UOS). Hydrogen bond-participating residues were identified in the wild type structures using the HbondMetric in RosettaScripts, with settings configured to exclude self-self hydrogen bonds. For each protein, residues engaged in at least one, two, or three hydrogen bonds were independently selected for mutation in three separate experiments. Then, systematic substitutions of these hydrogen bond-participating residues were performed, replacing each with a non-polar, hydrophobic amino acid (A, V, L, I, F, M, or G). These sequence variants were generated using ProteinMPNN [36], resulting in ten mutated sequences per protein system. Structural models of these variants were subsequently predicted with the LocalColabFold implementation of AF2 [26]. All resulting structural models, including the mutated variants and the unmodified reference structures, were evaluated using the ELEN model and its variants. Per-residue ELEN scores for the mutated models were then compared to the scores obtained from the corresponding reference structures to assess the impact of polar-to-hydrophobic mutations on model quality predictions.

*Benchmarking ELEN on challenging AF2 targets*

To systematically assess the performance of ELEN across a range of protein structural classes and error types, we benchmarked the method using the same dataset presented in "The power and pitfalls of AF2 for structure prediction beyond rigid globular proteins" [37]. This dataset comprises twelve challenging protein

targets that represent a diverse set of structural classes—including flexible, multidomain, and non-globular proteins—for which AF2 predictions have been shown to deviate substantially from experimentally determined structures. Both AF2 models and corresponding crystal structures were obtained directly from the supplementary materials provided by the original publication. The AF2 models were then scored using the full ELEN model and superimposed onto their respective crystal structure for direct comparison.
For each protein target, per-residue quality scores were extracted from the B-factor columns of the AF2 or ELEN PDB files, respectively. Error regions within each model were defined either manually, based on visual inspection of superimposed structures in PyMOL, or by applying a residue-level lDDT threshold (<0.6) to identify structurally inaccurate segments (Table 7). For both, ELEN and plDDT, mean scores were computed across all residues as well as specifically within the identified error regions. To assess the ability of each metric to detect local structural flaws, we calculated the overlap and Jaccard index between the set of error residues defined by lDDT and those flagged by ELEN or plDDT using established score cutoffs. These metrics, along with target annotations and error types, were summarized in a comparative table to enable systematic evaluation of the sensitivity and specificity of ELEN versus plDDT across the benchmark set.

*Assessing ELEN for the detection of enhanced loop design*

To engineer variants of human duodenal cytochrome B with a redesigned active enhanced loop, we applied a fold-conditioned design workflow in which backbone flexibility was restricted to residues 180–190 while the rest of the scaffold remained fixed. Structure generation was performed using RFdiffusion with scaffold guidance, followed by RosettaRelax energy minimization that restrained heme-binding residues to their native conformations. Sequence diversification was achieved using SolubleMPNN, generating multiple variants per structure while fixing catalytic and substrate/cofactor-binding residues identified as essential for activity. After further energy minimization, designs were iteratively filtered and refined over three cycles, with quality assessment at each step: ESMFold predictions were used to filter based on plDDT and backbone RMSD, and AF3 models were selected using pTM score thresholds. Final candidates were subjected to additional RosettaRelax refinement and filtered by total energy score, with restraint settings consistent across all rounds. For comparison, a control protocol using the native ironzyme structure and an equivalent sequence design and filtering strategy (but without targeted loop flexibility) was employed. Throughout both protocols, the set of fixed residues for each step was chosen based on structural and functional annotation to ensure the preservation of catalytic function and cofactor binding.

*ELEN detects disordered and variable regions in de novo designed enzymes*

Eight computational models, as well as the experimental crystal structure for the *de novo* designed Kemp eliminases, were obtained by [38]. The computational models were structurally aligned to the crystal structure using PyMOL's align function to enable residue-level comparison. Regions lacking electron density in the crystal structure were identified from the corresponding PDB file and annotated for comparison with ELEN predictions.

*Interpreting ELEN's internal representations with PCA*

To assess which geometric and physicochemical features were captured by ELEN and its variants, we analyzed the activations of the penultimate layer of the network's multilayer perceptron on a per-residue basis for each input protein loop. The dataset was prepared as described in Section 3.3, but restricted to one month of data to limit the number of datapoints (253 protein models). For each residue, relevant physicochemical features were computed and associated with the corresponding activation vectors. Dimensionality reduction was performed independently using PCA and Uniform Manifold Approximation and Projection to project the high-dimensional activation data into two dimensions. The resulting two-dimensional embeddings were visualized, with individual residues colored according to their respective physicochemical or geometric features, in order to qualitatively evaluate the feature sensitivity and representational structure learned by the network.

### Supplementary Results

#### *The ELEN loop dataset: Coverage and diversity*

***Supplementary Table S1.*** *Descriptive statistics for per-residue quality metrics (lDDT, CAD-score, RMSD) computed across all extracted loop pockets (n ≈ 1.5 million) in the ELEN loop dataset.*

| Statistic | lDDT score | CAD-score | RMSD (Å) |
|---|---|---|---|
| Mean | 0.901 | 0.835 | 1.61 |
| Standard deviation | 0.109 | 0.118 | 2.89 |
| Median | 0.938 | 0.870 | 0.83 |
| 25th percentile | 0.880 | 0.803 | 0.47 |
| 75th percentile | 0.964 | 0.909 | 1.82 |
| Skewness | -3.15 | -2.40 | 8.09 |
| Kurtosis | 13.1 | 7.81 | 103 |

#### *Detailed feature selection and ablation study*

To investigate the impact of additional input features on the performance of the ELEN model, we tested the concatenation of various per-residue features either to the initial atom-level embeddings or to the learned residue-level representations (Figure 1C). Predictive performance was quantified by calculating Pearson correlations and MAE between model predictions and ground truth labels (lDDT, CAD-Score, RMSD) on the test dataset during training. Supplementary Figure S1 displays the average Pearson correlation R, while Supplementary Table S2 presents detailed individual correlations and MAE across all three labels.
At the atom-level, continuous per-residue features significantly improved model predictions. Among these, Rosetta Energies yielded the highest correlation, with an average Pearson's R of 0.324 ± 0.011 for the lDDT metric. SASA and SAP-score also significantly improved predictions compared to baseline models that incorporated no additional features (R = 0.218 ± 0.012). Combining all continuous features at the atom-level further increased predictive performance (average R = 0.3702), indicating complementary information derived from multiple features. Categorical features, particularly sequence information, provided noticeable improvements over the baseline (R = 0.262), while secondary structure contributed moderate enhancements (R = 0.224). Additionally, we tested alternative initial atom-level encodings, including the one-hot element encoding proposed previously [43], but observed no improvement in predictive performance compared to our original encoding scheme. At the residue-level, continuous features similarly increased model predictions, with Rosetta Energies (R = 0.324 ± 0.011) and SASA (R = 0.295 ± 0.022) being again the most effective ones. The combination of all continuous features achieved an average R of 0.349 ± 0.022, clearly surpassing baseline (R = 0.218 ± 0.012). Categorical residue-level features provided limited overall predictive benefit, with Sequence (R = 0.262 ± 0.015) slightly outperforming secondary structure (R = 0.224 ± 0.012). Counting hydrogen bonds yielded modest improvements both at atom-level (R = 0.215 ± 0.012) and residue-level (R = 0.239 ± 0.022), underscoring their limited but consistent predictive utility.

Incorporation of sequence embeddings derived from LLMs further enhanced predictive performance. We systematically evaluated various sequence embedding models, including multiple ESM-2 variants and several sizes of SaProt models. Among these, the SaProt model version SaProt_650M_PDB demonstrated superior predictive accuracy, providing substantial improvements at the atom-level ($R = 0.397 \pm 0.116$) and notable enhancements at the residue-level ($R = 0.440 \pm 0.018$). Additionally, various per-atom features (e.g., number of hydrogen bonds per atom, atomic B-factors, and charges) were tested but ultimately excluded due to computational complexity and minimal additional predictive performance compared to per-residue descriptors. In conclusion, carefully selected continuous descriptors, particularly advanced embeddings such as SaProt LLM, significantly enhance ELEN's ability to predict structural quality, suggesting their integration as best practices for future model optimization.

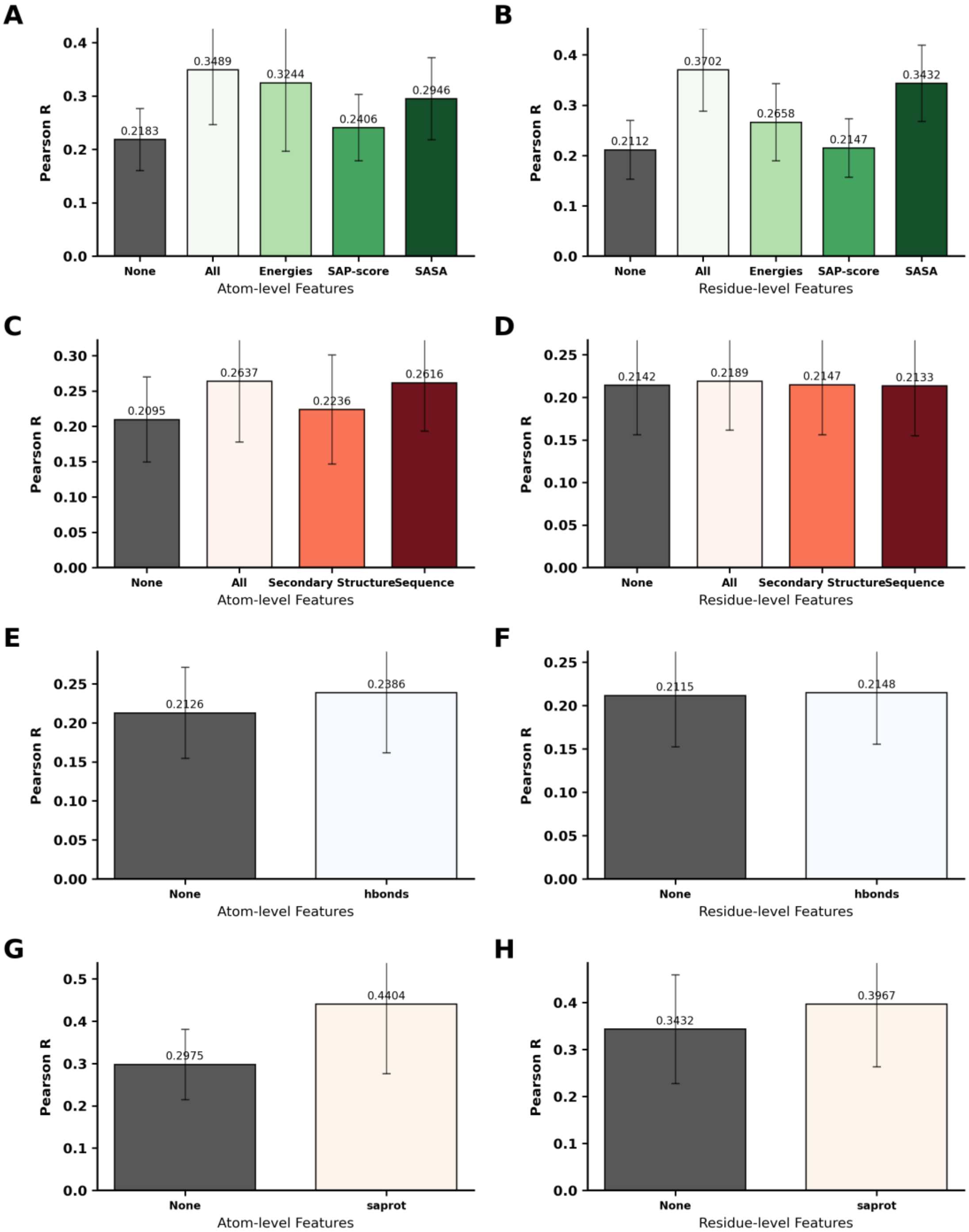


***Supplementary Figure S1. Influence of atom-level and residue-level features on model performance.*** *Pearson's correlation coefficient (R) between predicted and observed quality metrics (CAD-score, lDDT, and RMSD) is shown for models incorporating different combinations of atom-level (left, panels A, C, E, G) and residue-level (right, panels B, D, F, H) features. Features are categorized as continuous (Energies, SAP-score, SASA), categorical (secondary structure, sequence), count-based (H-bonds) or sequence embeddings (SaProt). Bars represent average correlation across metrics, with error bars depicting standard deviation. The "None" category indicates models without additional input features, serving as baseline performance. "All" denotes averaged performance across all feature categories within each layer and feature type.*

***Supplementary Table S2. Summary of model performance using various atom-level and residue-level features.*** *For each feature set, Pearson's correlation coefficient (R) and MAE are reported across three structural quality metrics: lDDT, CAD-score, and RMSD. Means (***μ***), standard deviations (***σ***), and sample sizes (N) are shown for each feature group. Feature types include continuous variables (Energies, SAP-score, SASA), categorical variables (sequence, secondary structure), count-based features (number of H-bonds), and sequence embeddings derived from the SaProt LLM. The category "All features" indicates models trained with all features of the corresponding subcategory combined. "AVG" denotes the mean across all three labels.*

| | | | | | **lDDT** | | | | **CAD-score** | | | | **RMSD** | | | |
|---|---|---|---|---|---|---|---|---|---|---|---|---|---|---|---|---|
| | | | | | **R** | | **MAE** | | **R** | | **MAE** | | **R** | | **MAE** | |
| **Layer** | **Feature** | **Type** | **N** | **AVG** | **μ** | **σ** | **μ** | **σ** | **μ** | **σ** | **μ** | **σ** | **μ** | **σ** | **μ** | **σ** |
| Atom | None | cont. | 41 | 0.211 | 0.267 | 0.014 | 0.059 | 0.003 | 0.216 | 0.013 | 0.074 | 0.003 | 0.151 | 0.021 | 0.935 | 0.268 |
| Atom | All features | cont. | 35 | 0.370 | 0.452 | 0.006 | 0.054 | 0.003 | 0.371 | 0.008 | 0.070 | 0.003 | 0.288 | 0.016 | 0.934 | 0.322 |
| Atom | Energies | cont. | 40 | 0.266 | 0.331 | 0.013 | 0.056 | 0.002 | 0.285 | 0.014 | 0.071 | 0.002 | 0.181 | 0.023 | 0.971 | 0.331 |
| Atom | SAP-Score | cont. | 41 | 0.215 | 0.272 | 0.013 | 0.059 | 0.002 | 0.217 | 0.013 | 0.074 | 0.002 | 0.156 | 0.017 | 0.946 | 0.276 |
| Atom | SASA | cont. | 43 | 0.343 | 0.424 | 0.009 | 0.056 | 0.003 | 0.332 | 0.008 | 0.072 | 0.002 | 0.274 | 0.018 | 1.007 | 0.300 |
| Atom | None | cat. | 56 | 0.214 | 0.271 | 0.014 | 0.060 | 0.005 | 0.217 | 0.014 | 0.074 | 0.002 | 0.154 | 0.019 | 0.962 | 0.225 |
| Atom | All features | cat. | 21 | 0.264 | 0.347 | 0.021 | 0.059 | 0.004 | 0.270 | 0.020 | 0.075 | 0.004 | 0.175 | 0.036 | 1.029 | 0.466 |
| Atom | Secondary structure | cat. | 38 | 0.215 | 0.272 | 0.014 | 0.059 | 0.004 | 0.217 | 0.016 | 0.074 | 0.002 | 0.155 | 0.022 | 0.923 | 0.244 |
| Atom | Sequence | cat. | 56 | 0.213 | 0.270 | 0.014 | 0.059 | 0.003 | 0.217 | 0.015 | 0.074 | 0.002 | 0.153 | 0.021 | 0.943 | 0.237 |
| Atom | None | counts | 47 | 0.212 | 0.269 | 0.012 | 0.059 | 0.002 | 0.215 | 0.012 | 0.074 | 0.002 | 0.151 | 0.028 | 0.909 | 0.195 |
| Atom | Number of H-bonds | counts | 48 | 0.215 | 0.273 | 0.012 | 0.059 | 0.003 | 0.218 | 0.015 | 0.074 | 0.002 | 0.154 | 0.017 | 0.938 | 0.269 |
| Atom | None | cont. | 75 | 0.343 | 0.444 | 0.136 | 0.056 | 0.010 | 0.369 | 0.111 | 0.070 | 0.008 | 0.217 | 0.048 | 1.271 | 0.450 |
| Atom | SaProt LLM | cont. | 88 | 0.397 | 0.513 | 0.116 | 0.053 | 0.009 | 0.427 | 0.097 | 0.068 | 0.008 | 0.251 | 0.053 | 1.133 | 0.426 |
| Residue | None | cont. | 23 | 0.218 | 0.276 | 0.012 | 0.059 | 0.004 | 0.219 | 0.016 | 0.074 | 0.002 | 0.159 | 0.023 | 0.971 | 0.317 |
| Residue | All features | cont. | 20 | 0.349 | 0.367 | 0.022 | 0.055 | 0.004 | 0.441 | 0.020 | 0.071 | 0.003 | 0.238 | 0.023 | 0.901 | 0.260 |
| Residue | Energies | cont. | 15 | 0.324 | 0.427 | 0.011 | 0.053 | 0.002 | 0.365 | 0.009 | 0.069 | 0.002 | 0.181 | 0.042 | 1.020 | 0.310 |
| Residue | SAP-Score | cont. | 24 | 0.241 | 0.307 | 0.021 | 0.061 | 0.003 | 0.231 | 0.019 | 0.076 | 0.005 | 0.184 | 0.028 | 1.105 | 0.347 |
| Residue | SASA | cont. | 28 | 0.295 | 0.374 | 0.022 | 0.061 | 0.004 | 0.289 | 0.022 | 0.075 | 0.003 | 0.220 | 0.020 | 0.941 | 0.291 |
| Residue | None | cat. | 24 | 0.210 | 0.268 | 0.009 | 0.058 | 0.003 | 0.213 | 0.013 | 0.074 | 0.002 | 0.148 | 0.017 | 0.928 | 0.305 |

| | | | | | | | | | | | | | | | | |
|---|---|---|---|---|---|---|---|---|---|---|---|---|---|---|---|---|
| Residue | All features | cat. | 50 | 0.219 | 0.275 | 0.014 | 0.059 | 0.003 | 0.221 | 0.015 | 0.075 | 0.002 | 0.161 | 0.019 | 0.947 | 0.331 |
| Residue | Secondary structure | cat. | 21 | 0.224 | 0.301 | 0.012 | 0.061 | 0.004 | 0.224 | 0.015 | 0.078 | 0.003 | 0.146 | 0.030 | 0.962 | 0.223 |
| Residue | Sequence | cat. | 36 | 0.262 | 0.326 | 0.014 | 0.057 | 0.003 | 0.269 | 0.014 | 0.072 | 0.002 | 0.190 | 0.030 | 1.013 | 0.311 |
| Residue | None | counts | 47 | 0.213 | 0.270 | 0.012 | 0.059 | 0.002 | 0.215 | 0.016 | 0.074 | 0.002 | 0.153 | 0.015 | 0.993 | 0.372 |
| Residue | Number of H-bonds | counts | 37 | 0.239 | 0.312 | 0.022 | 0.060 | 0.004 | 0.246 | 0.020 | 0.075 | 0.002 | 0.158 | 0.020 | 0.897 | 0.144 |
| Residue | None | cont. | 78 | 0.298 | 0.372 | 0.110 | 0.058 | 0.011 | 0.312 | 0.094 | 0.072 | 0.009 | 0.208 | 0.058 | 1.242 | 0.466 |
| Residue | SaProt LLM | cont. | 85 | 0.440 | 0.581 | 0.018 | 0.051 | 0.007 | 0.480 | 0.016 | 0.066 | 0.006 | 0.260 | 0.034 | 1.154 | 0.416 |
| Atom | None | cont. | 93 | 0.215 | 0.272 | 0.012 | 0.059 | 0.003 | 0.217 | 0.014 | 0.074 | 0.002 | 0.155 | 0.018 | 0.922 | 0.259 |
| Atom | b-factor | cont. | 32 | 0.277 | 0.349 | 0.017 | 0.056 | 0.002 | 0.269 | 0.017 | 0.073 | 0.002 | 0.212 | 0.017 | 0.929 | 0.219 |
| Atom | All features | cont. | 32 | 0.279 | 0.350 | 0.015 | 0.056 | 0.002 | 0.270 | 0.011 | 0.073 | 0.002 | 0.216 | 0.020 | 0.885 | 0.166 |
| Atom | charges | cont. | 35 | 0.222 | 0.280 | 0.014 | 0.060 | 0.004 | 0.223 | 0.016 | 0.074 | 0.003 | 0.163 | 0.016 | 0.981 | 0.279 |
| Atom | H-bonds | counts | 33 | 0.217 | 0.279 | 0.013 | 0.060 | 0.003 | 0.220 | 0.012 | 0.075 | 0.002 | 0.152 | 0.017 | 1.030 | 0.315 |

*Supplementary figures and tables for the study of the structural and geometric awareness of ELEN*

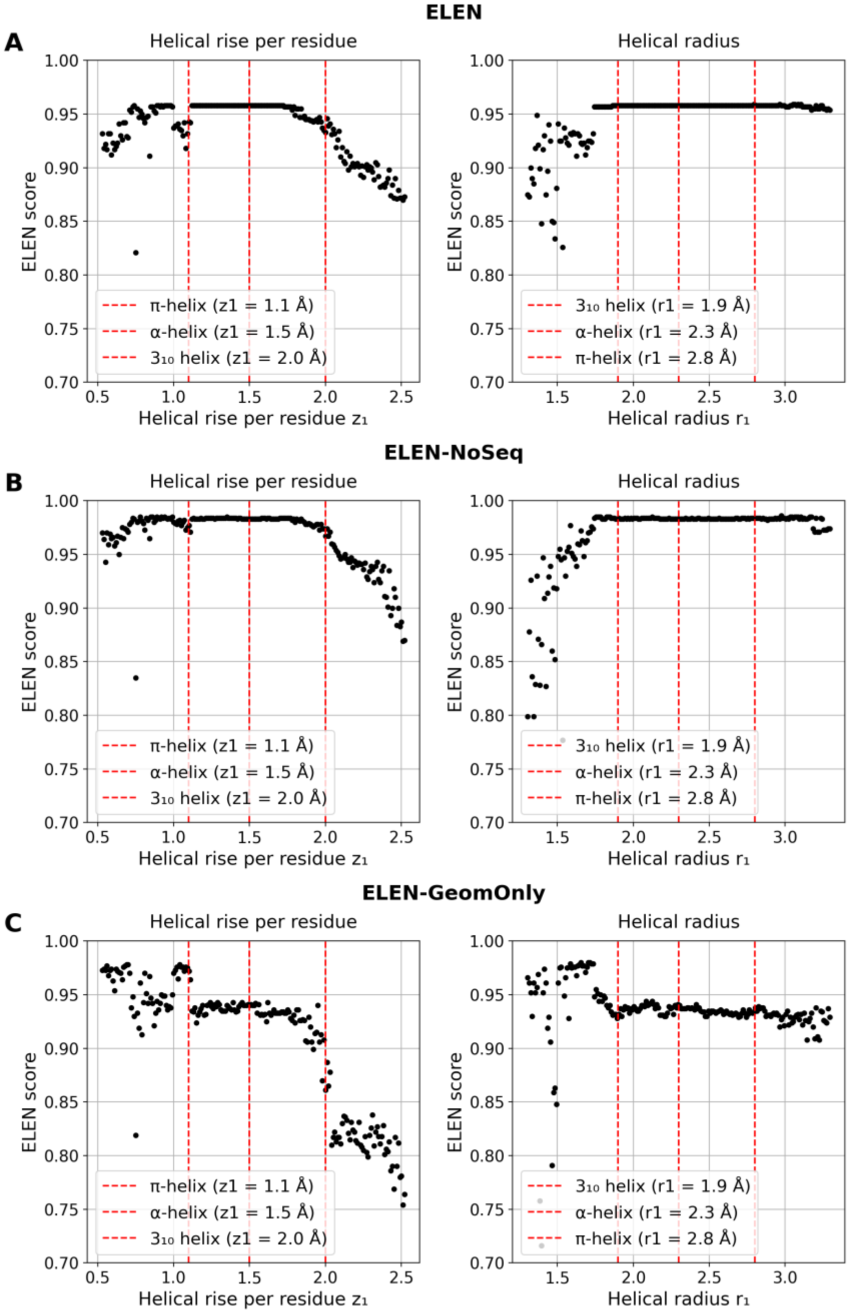


***Supplementary Figure S2. Assessment of the geometric and structural sensitivity of ELEN and its variants.*** *For each ELEN variant (A: ELEN, B: ELEN-NoSeq, C: ELEN-GeomOnly), the averaged per-residue ELEN score is plotted as a function of helical rise per residue $z_1$ (left) and helical radius $r_1$ (right) for a series of idealized 25-residue helices generated using systematically varied helical parameters. Vertical red dashed lines indicate canonical values for π-helix, α-helix, and $3_{10}$-helix geometries.*

***Supplementary Table S3:*** *Mean ELEN score at canonical helix parameters for each model variant. Scores are reported for π-helix, α-helix, and $3_{10}$-helix rises ($z_1$) and radii ($r_1$). Higher scores indicate closer agreement with native-like geometry.*

| | **helical rise per residue $z_1$** | | | **helical radius $r_1$** | | |
|---|---|---|---|---|---|---|
| **Model variant** | **π-helix (1.1 Å)** | **α-helix (1.5 Å)** | **$3_{10}$-helix (2.0 Å)** | **π-helix (2.8 Å)** | **α-helix (2.3 Å)** | **$3_{10}$-helix (1.9 Å)** |
| ELEN | 0.943 | 0.958 | 0.933 | 0.958 | 0.958 | 0.958 |
| ELEN-NoSeq | 0.977 | 0.983 | 0.967 | 0.984 | 0.983 | 0.984 |
| ELEN-GeomOnly | 0.972 | 0.94 | 0.861 | 0.938 | 0.94 | 0.932 |

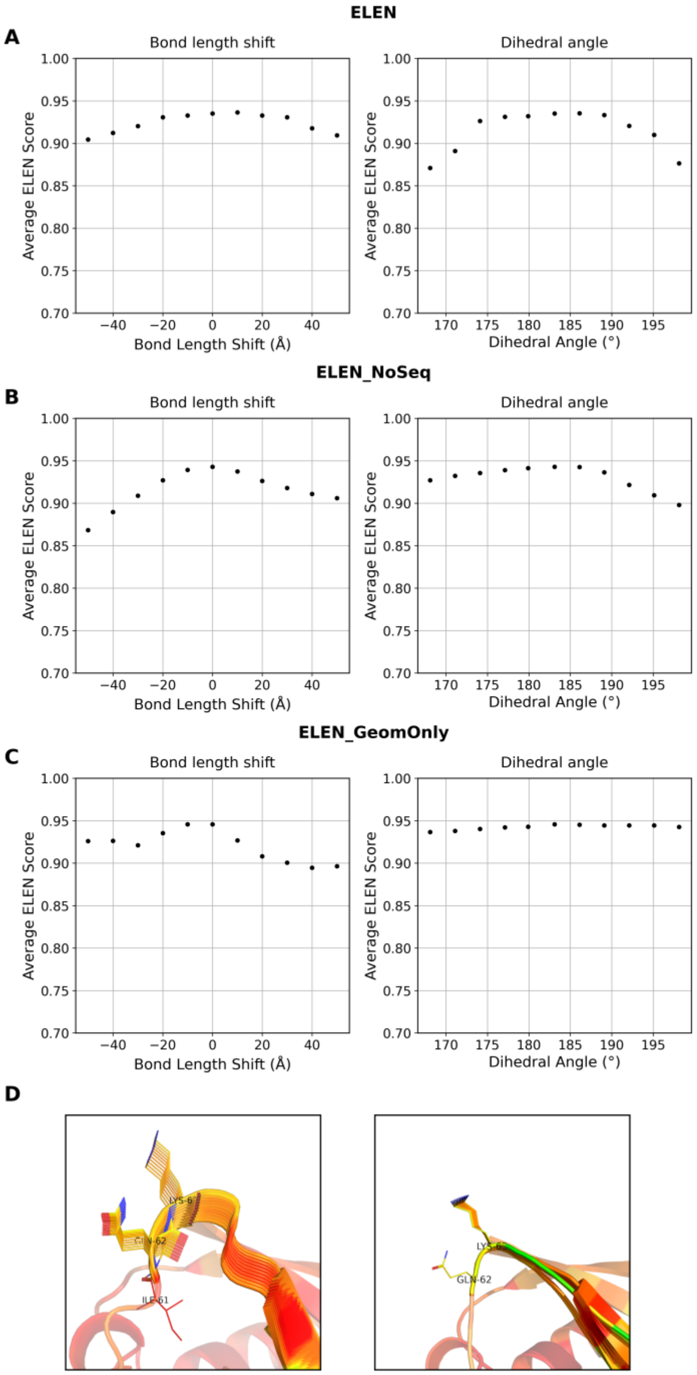


***Supplementary Figure S3. Sensitivity of ELEN model variants to systematic backbone perturbations in ubiquitin (PDB ID: 1UBQ).*** *(A–C) Average ELEN scores for residues 61–63 as a function of backbone bond length shift (left) and peptide bond dihedral angle ω (right) for (A) the full ELEN model (A), (B) ELEN_NoSeq (B), and ELEN_GeomOnly (C). (D) Structural superpositions illustrating the effects of bond length (left) and dihedral angle (right) perturbations at residue 62. Color gradient represents the ELEN score from red (highest) through yellow to green (lowest), where red indicates the best score.*

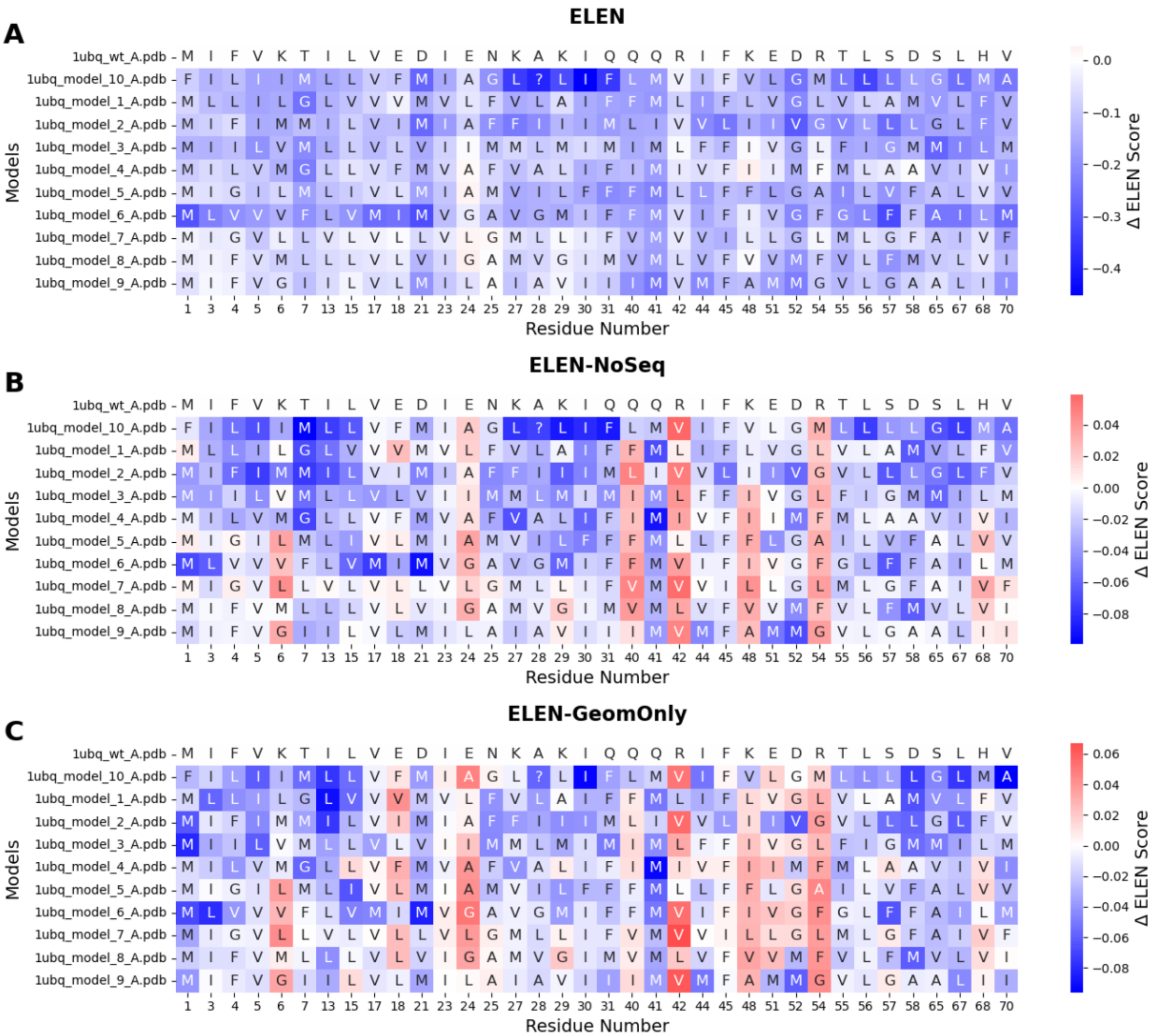


***Supplementary Figure S4. Impact of polar-to-hydrophobic mutations on per-residue ELEN scores in ubiquitin.*** *Heatmaps illustrate the impact of polar-to-hydrophobic mutations (at residues forming at least two hydrogen bonds) on per-residue ELEN scores in ubiquitin (PDB ID: 1UBQ). The changes (ΔELEN) relative to the wild-type structure are shown for (A) the full ELEN model, (B) the sequence-agnostic ELEN-NoSeq variant, and (C) the geometry-only ELEN-GeomOnly variant. Blue indicates a reduction in predicted structural quality, whereas red denotes an improvement.*

***Supplementary Table S4.*** *Summary of mean ELEN scores and their differences (ΔELEN) between wild-type and mutated structural models across varying thresholds of hydrogen bond participation (≥1, ≥2, ≥3 H-bonds) for ubiquitin (PDB ID: 1UBQ) and the four-helix bundle (PDB ID: 4UOS). Results are reported for the full ELEN model and two variant models ("NoSeq" and "GeomOnly"). Means were computed over per-residue values as well as across the 10 independently mutated sequences. Standard deviations (SD) reflect variability across these 10 sequence replicates.*

| PDB ID | H-bond cutoff | Model variant | $\mu_{Orig}$ | $\mu_{Mut}$ | $\sigma_{Mut}$ | ΔELEN ($\mu + \sigma$) |
|---|---|---|---|---|---|---|
| 1UBQ | 1 | ELEN | 0.881 | 0.612 | 0.095 | -0.269 ± 0.096 |

| | | | | | | |
|---|---|---|---|---|---|---|
| 1UBQ | 1 | NoSeq | 0.956 | 0.884 | 0.060 | -0.071 ± 0.062 |
| 1UBQ | 1 | GeomOnly | 0.952 | 0.876 | 0.058 | -0.075 ± 0.058 |
| 1UBQ | 2 | ELEN | 0.887 | 0.766 | 0.076 | -0.121 ± 0.073 |
| 1UBQ | 2 | NoSeq | 0.957 | 0.941 | 0.026 | -0.017 ± 0.029 |
| 1UBQ | 2 | GeomOnly | 0.953 | 0.940 | 0.024 | -0.013 ± 0.029 |
| 1UBQ | 3 | ELEN | 0.914 | 0.859 | 0.029 | -0.055 ± 0.023 |
| 1UBQ | 3 | NoSeq | 0.969 | 0.957 | 0.014 | -0.012 ± 0.010 |
| 1UBQ | 3 | GeomOnly | 0.967 | 0.962 | 0.009 | -0.004 ± 0.008 |
| 4UOS | 1 | ELEN | 0.848 | 0.671 | 0.087 | -0.177 ± 0.090 |
| 4UOS | 1 | NoSeq | 0.943 | 0.902 | 0.062 | -0.041 ± 0.067 |
| 4UOS | 1 | GeomOnly | 0.929 | 0.854 | 0.062 | -0.075 ± 0.067 |
| 4UOS | 2 | ELEN | 0.858 | 0.721 | 0.074 | -0.137 ± 0.076 |
| 4UOS | 2 | NoSeq | 0.947 | 0.901 | 0.052 | -0.046 ± 0.052 |
| 4UOS | 2 | GeomOnly | 0.933 | 0.865 | 0.052 | -0.068 ± 0.056 |
| 4UOS | 3 | ELEN | 0.840 | 0.838 | 0.034 | -0.002 ± 0.025 |
| 4UOS | 3 | NoSeq | 0.940 | 0.952 | 0.020 | 0.012 ± 0.024 |
| 4UOS | 3 | GeomOnly | 0.921 | 0.941 | 0.023 | 0.020 ± 0.026 |

#### *Supplementary material for section Assessing ELEN in AF2 failure cases and advanced design scenarios*

***Supplementary Table S5. Performance of ELEN and plDDT on challenging protein structure prediction targets.*** *Summary of ELEN and AF2's plDDT scores on twelve proteins previously identified as difficult cases for structure prediction by AF2. Target types, main error classes, and error region detection methods (manual curation or lDDT thresholding) are indicated for each protein. For both ELEN and plDDT, mean scores are shown across all residues ($\mu_{ELEN}$) and for residues in the error region ($\mu_{Error}$). Overlap and Jaccard indices quantify the agreement between residues identified as erroneous by lDDT and those flagged by ELEN or plDDT (using defined thresholds). "Flaw?" columns indicate whether each method successfully detected the main error region.*

| PDB ID | Type | Main Error | Error region detection | $\mu_{lDDT}$ | $\mu_{ELEN}$ | $\mu_{error}$ | Overlap (%) | Jaccard (%) | ELEN flaw? | plDDT (avg) | plDDT (err avg) | Overlap (%) | Jaccard (%) | plDDT flaw? |
|---|---|---|---|---|---|---|---|---|---|---|---|---|---|---|
| 1S4T | Peptide | misfold | manually | 0.399 | 0.723 | 0.723 | 85.7 | 85.7 | Yes | 88.08 | 88.08 | 0.0 | 0.0 | No |
| 2KQP | Globular | misfold | manually | 0.666 | 0.421 | 0.421 | 100.0 | 100.0 | Yes | 48.14 | 48.14 | 100.0 | 100.0 | Yes |
| 2Y4Q | Cofactors/ligands | conformational change | lDDT | 0.634 | 0.709 | 0.644 | 100.0 | 50.0 | Yes | 66.19 | 61.67 | 89.7 | 45.6 | Yes |
| 6OFS | Enzymes | conformational change | manually | 0.803 | 0.885 | 0.878 | 4.7 | 4.7 | No | 93.01 | 91.81 | 1.2 | 1.2 | No |
| 3T5O | Extended/repeats | domain orientation | manually | 0.757 | 0.825 | 0.828 | 70.5 | 31.3 | No | 81.50 | 82.14 | 58.0 | 46.1 | No |
| 7W9W | Membrane proteins | local misaligment | lDDT | 0.713 | 0.787 | 0.604 | 86.5 | 59.3 | Yes | 83.81 | 66.90 | 43.2 | 42.1 | Yes |
| 1WVK | IDPs/IDRs | misfold | lDDT | 0.470 | 0.836 | 0.832 | 20.9 | 20.6 | Partly | 78.89 | 78.59 | 29.9 | 29.9 | Partly |
| 1TF2 | Conformational change | conformational change | lDDT | 0.803 | 0.871 | 0.793 | 43.8 | 23.5 | Partly | 89.62 | 76.76 | 30.14 | 22.9 | Partly |
| 4ORW | Mutations | local misaligment | lDDT | 0.832 | 0.834 | 0.740 | 70.6 | 26.1 | Yes | 93.08 | 81.37 | 29.4 | 29.4 | No |
| 6W90 | De novo design | local misaligment | lDDT | 0.638 | 0.688 | 0.699 | 73.3 | 34.4 | Yes | 89.38 | 84.50 | 4.4 | 4.3 | No |
| 3RJQ | Protein complex | domain orientation | manually | 0.811 | 0.816 | 0.776 | 81.4 | 32.4 | No | 89.39 | 84.68 | 51.2 | 41.5 | No |
| 6TUB | Oligomer/amyloid | misfold | lDDT | 0.382 | 0.536 | 0.536 | 100.0 | 94.1 | Yes | 36.62 | 36.62 | 100.0 | 94.1 | Yes |

**Supplementary Table S6.** ***Statistical summary of model quality and structural metrics for original and redesigned protein variants across different regions.*** *For each group (original and redesign) and region (loop: residues 180–190; rest: residues 1–179 and 191–225; whole protein), the table reports the number of models (N), mean (μ), standard deviation (σ), median, minimum, and maximum values for ELEN (predicted model quality), plDDT and backbone RMSD (Å) relative to the original model. p-values were calculated using Welch's t-test (two-sided, unequal variance) to assess the significance of differences between original and redesigned groups for each metric.*

| | | | ELEN | | | | | plDDT | | | | | RMSD [Å] | | | |
|---|---|---|---|---|---|---|---|---|---|---|---|---|---|---|---|---|
| Region | Model | N | μ + σ | Median | Min | Max | p-value | μ + σ | Median | Min | Max | p-value | μ + σ | Median | Min | Max |
| loop (180-190) | original | 161 | 0.812 ± 0.038 | 0.819 | 0.676 | 0.878 | 1.55E-06 | 70.31 ± 8.10 | 70.93 | 50.23 | 83.96 | 5.75E-17 | 2.34±1.24 | 1.91 | 0.94 | 7.86 |
| | redesign | 52 | 0.840 ± 0.032 | 0.851 | 0.761 | 0.896 | 1.55E-06 | 82.43 ± 7.23 | 84.39 | 65.67 | 91.74 | 5.75E-17 | 5.64±0.80 | 5.68 | 3.78 | 7.06 |
| rest (1-179, 191-225) | original | 161 | 0.881 ± 0.004 | 0.881 | 0.871 | 0.892 | 1.71E-48 | 87.62 ± 1.60 | 87.65 | 83.92 | 91.25 | 2.91E-29 | 1.17±0.07 | 1.16 | 1.03 | 1.48 |
| | redesign | 52 | 0.896 ± 0.003 | 0.896 | 0.884 | 0.902 | 1.71E-48 | 90.83 ± 1.21 | 90.99 | 88.51 | 93.48 | 2.91E-29 | 1.15±0.05 | 1.15 | 1.05 | 1.48 |
| whole protein | original | 161 | 0.878 ± 0.005 | 0.878 | 0.862 | 0.891 | 9.78E-48 | 86.78 ± 1.71 | 86.83 | 82.71 | 90.75 | 2.31E-31 | 1.23±0.11 | 1.20 | 1.05 | 1.60 |
| | redesign | 52 | 0.893 ± 0.004 | 0.893 | 0.883 | 0.899 | 9.78E-48 | 90.42 ± 1.29 | 90.65 | 87.97 | 93.10 | 2.31E-31 | 1.37±0.07 | 1.38 | 1.19 | 1.56 |

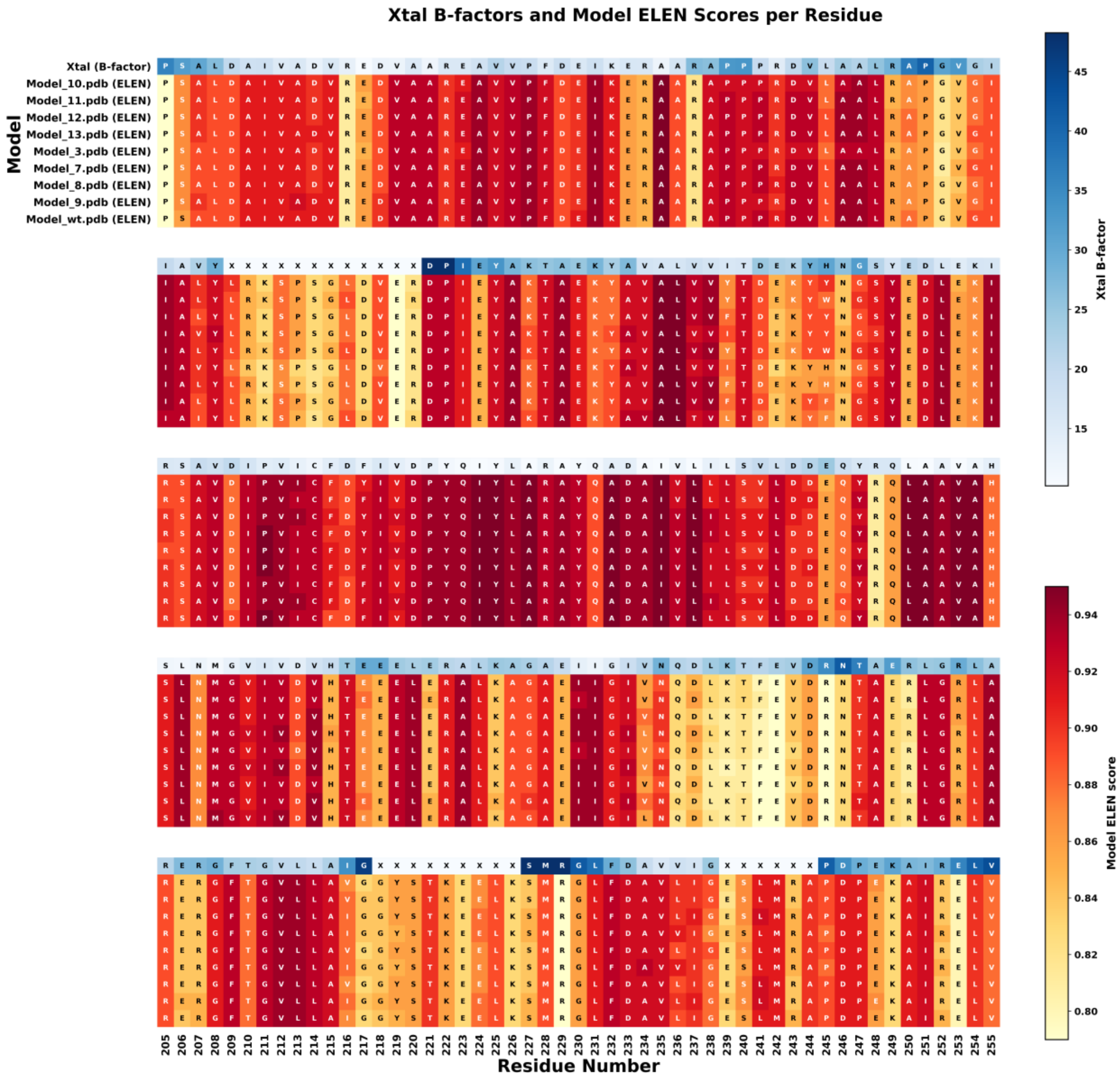


***Supplementary Figure S5. Per-residue quality heatmap comparing experimental B-factors and ELEN scores for Kemp eliminase designs.*** *The top row shows the crystal structure colored by experimental B-factors, with the corresponding amino acid sequence annotated for each residue. Lower rows represent individual computational models, colored by per-residue ELEN scores, with sequence letters overlaid. Residues assigned as 'X' indicate positions lacking clear electron density in the crystal structure. Regions of elevated B-factor and high ELEN score consistently highlight variable or disordered segments, enabling direct comparison of experimental flexibility and predicted model uncertainty across the ensemble. Color bars indicate the range and scale for B-factors and ELEN scores.*

***Supplementary Table S7. Summary statistics of ELEN model quality scores for all, resolved, and disordered (missing density) regions in de novo designed Kemp eliminase models.*** *For each computational model, the mean (μ) and standard deviation (σ) of the ELEN score are shown for all residues, for regions resolved in the crystal structure (i.e., present in electron density), and for each of three regions absent from the crystal structure ("disordered xtal"). Δ indicates the difference between the mean ELEN score in resolved regions and each missing region. Averages across all models are shown in the last row. The crystal structure itself was also evaluated with the ELEN model, and these results, along with the corresponding experimental B-factors, are included for comparison. Regions absent from the crystal structure lack experimental B-factors and ELEN scores for the xtal reference.*

| | **All** | **Ordered** | **Disordered xtal region 1** | | **Disordered xtal region 2** | | **Disordered xtal region 3** | |
|---|---|---|---|---|---|---|---|---|
| **Model** | **μ + σ** | **μ + σ** | **μ + σ** | **Δ(Ord - Dis1)** | **μ + σ** | **Δ(Ord - Dis2)** | **μ + σ** | **Δ(Ord - Dis3)** |
| Model_10 | 0.894 ± 0.041 | 0.899 ± 0.040 | 0.850 ± 0.034 | 0.049 | 0.859 ± 0.024 | 0.040 | 0.877 ± 0.034 | 0.022 |
| Model_11 | 0.895 ± 0.041 | 0.899 ± 0.040 | 0.853 ± 0.033 | 0.047 | 0.860 ± 0.024 | 0.040 | 0.876 ± 0.038 | 0.023 |
| Model_12 | 0.895 ± 0.041 | 0.899 ± 0.040 | 0.850 ± 0.033 | 0.049 | 0.861 ± 0.025 | 0.039 | 0.878 ± 0.035 | 0.022 |
| Model_13 | 0.896 ± 0.041 | 0.899 ± 0.040 | 0.849 ± 0.032 | 0.051 | 0.860 ± 0.023 | 0.040 | 0.878 ± 0.032 | 0.022 |
| Model_3 | 0.895 ± 0.041 | 0.899 ± 0.040 | 0.852 ± 0.033 | 0.047 | 0.858 ± 0.024 | 0.040 | 0.878 ± 0.034 | 0.021 |
| Model_7 | 0.894 ± 0.043 | 0.899 ± 0.042 | 0.842 ± 0.032 | 0.057 | 0.859 ± 0.024 | 0.040 | 0.876 ± 0.034 | 0.023 |
| Model_8 | 0.894 ± 0.042 | 0.899 ± 0.041 | 0.845 ± 0.033 | 0.054 | 0.859 ± 0.024 | 0.039 | 0.877 ± 0.035 | 0.022 |
| Model_9 | 0.896 ± 0.041 | 0.899 ± 0.041 | 0.849 ± 0.033 | 0.050 | 0.861 ± 0.024 | 0.039 | 0.878 ± 0.034 | 0.022 |
| AVG Models | 0.895 ± 0.042 | 0.899 ± 0.041 | 0.849 ± 0.033 | 0.051 | 0.859 ± 0.024 | 0.039 | 0.877 ± 0.034 | 0.022 |
| xtal | 0.853 ± 0.096 | 0.853 ± 0.096 | - | | - | | - | |
| b-factor | 24.03 ± 8.94 | 24.03 ± 8.94 | - | | - | | - | |

*Supplementary material for section Interpreting ELEN's internal representations with PCA*

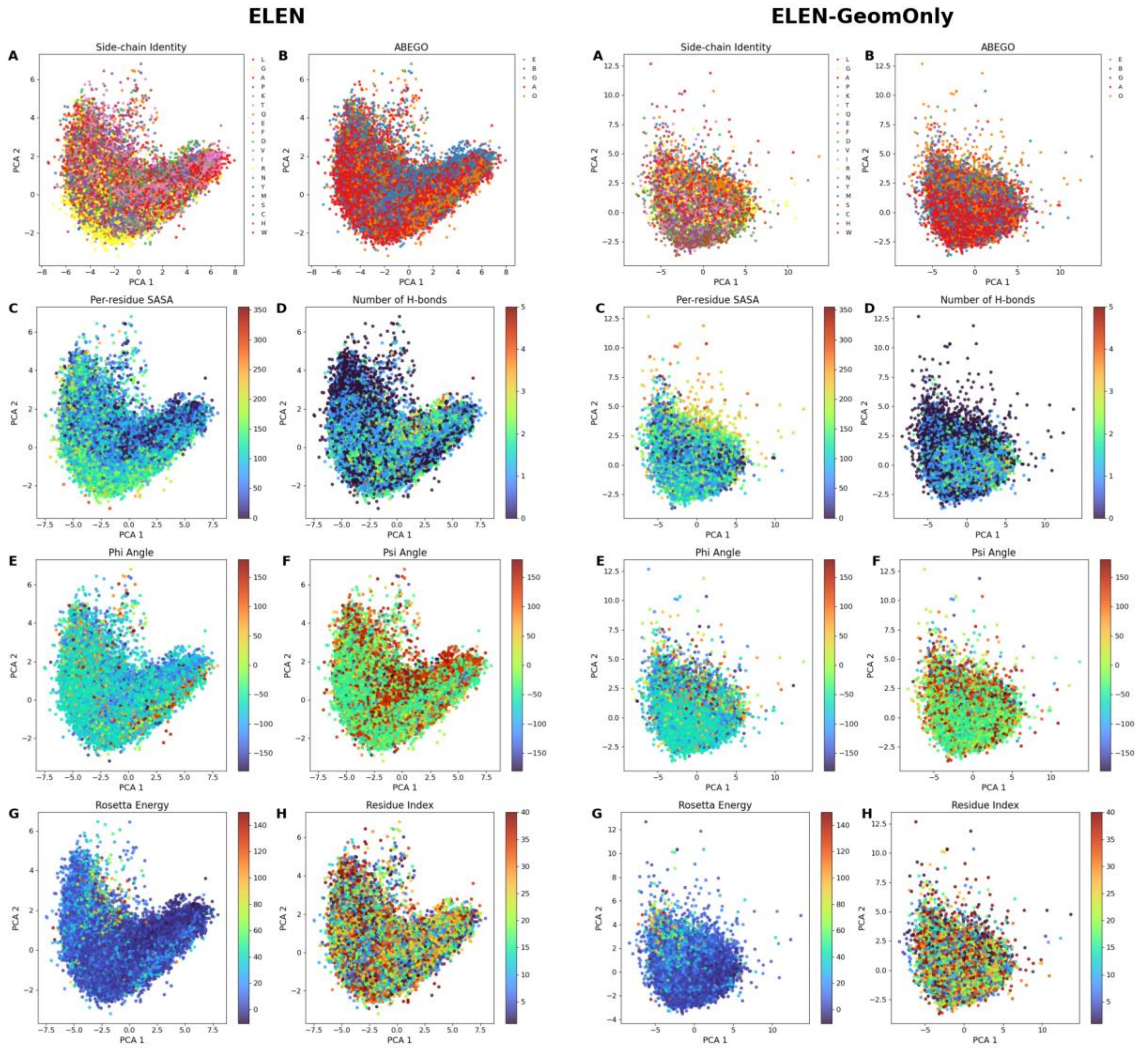


**Supplementary** ***Figure S6. Comparison of network activation feature sensitivity for ELEN (left) and ELEN-GeomOnly (right) models.*** *Scatter plots show the projection of penultimate-layer activations for all residues in the dataset onto the first two principal components (PCA1 and PCA2), with each point representing a single residue. PCA was performed on the penultimate layer activations for both the full ELEN model (left panels) and a variant trained with only geometric features (ELEN-GeomOnly, right panels). Points are colored according to (A) side-chain identity, (B) ABEGO torsion bin, (C) per-residue SASA, (D) number of hydrogen bonds, (E) phi angle, (F) psi angle, (G) Rosetta energy, and (H) residue index (as negative control). Legends or colorbars indicate the mapping of colors to feature values or categories.*
